# ProCogGraph: A Graph-Based Mapping of Cognate Ligand Domain Interactions

**DOI:** 10.1101/2024.08.08.607191

**Authors:** Matthew Crown, Matthew Bashton

**Affiliations:** Hub for Biotechnology in the Built Environment, Department of Applied Sciences, Faculty of Health and Life Sciences, Northumbria University, Newcastle upon Tyne, NE1 8ST, United Kingdom

## Abstract

**Motivation:** Mappings of domain-cognate ligand interactions can enhance our understanding of the core concepts of evolution and be used to aid docking and protein design. Since the last available cognate-ligand domain database was released, the PDB has grown significantly and new tools are available for measuring similarity and determining contacts.

**Results:** We present ProCogGraph, a graph database of cognate-ligand domain mappings in PDB structures. Building upon the work of the predecessor database, PROCOGNATE, we use data-driven approaches to develop thresholds and interaction modes. We explore new aspects of domain-cognate ligand interactions, including the chemical similarity of bound cognate ligands and how domain combinations influence cognate ligand binding. Finally, we use the graph to add specificity to partial EC IDs, showing that ProCogGraph can complete partial annotations systematically through assigned cognate ligands.

**Availability and Implementation:** The ProCogGraph pipeline, database and flat files are available at https://github.com/bashton-lab/ProCogGraph and https://doi.org/10.5281/zenodo.13165851.

**Supplementary information:** Supplementary data are available at *Bioinformatics* online.

## 1 Introduction

Protein domains are independently folding, stable, three-dimensional protein substructures first proposed in the 1970s (Rao and Rossmann 1973). Various databases and methods exist for assigning domains in experimental and computationally predicted protein structures, including CATH, SCOP2 and Pfam (Andreeva *et al*. 2020; Mistry *et al*. 2021; Waman *et al*. 2024).

In biochemistry, a ligand is a molecule which reversibly binds to a protein, typically as part of the overall function of a protein, and can be chemically and structurally diverse, including metal ions, small molecules, oligosaccharides and other polypeptides. When considering enzymes, an endogenous ligand can be considered a naturally occurring ligand, e.g. ATP or glucose, but could also include naturally occurring inhibitors of enzymes; for instance, in Aspartate Transcarbamoylase cytidine triphosphate (CTP) acts as an inhibitor to the reaction in a feedback loop, by binding to an allosteric site and inducing a conformational change in the overall structure of the enzyme (Lipscomb and Kantrowitz 2012). The term cognate ligand is used to describe the endogenous ligands a protein binds during normal function - and for enzymes describes the set of ligands (substrates, products, cofactors) involved in their normal function. In the case of experimentally determined structures, ligands are often present, which may be the expected in vivo cognate ligand, an exogenous analogue or inhibitor used to aid structural determination such as N-(phosphonoacetyl)-L-aspartate (PALA), a transition state analogue of the L-aspartate and carbamoyl phosphate substrates of the ATCase enzyme (Krause, Volz and Lipscomb 1987), or an artefact of crystallisation, e.g. a buffer component such as glycerol.

Many different databases exist to document protein-ligand binding interfaces. The PDBe-Graph (Nair *et al*. 2021) is a graph database of residue level annotations for PDB structures, including protein-ligand contacts and also includes mappings to protein domains in CATH, SCOP, Pfam and InterPro (Blum *et al*. 2021). Whilst the PDBe-graph contains a rich annotation, a limitation of this database is the lack of specific endpoints to extract protein domain specific interactions between a ligand and a protein. PDBsum is a web-server which provides summary information on PDB structures, including domain annotations and protein-ligand interactions, determined using HBPLUS, primarily focussed on schematics for visual analysis of proteins and provides a similarity measure between ligands in the structure and cognate substrates/products by comparing their maximum common substructure (MCS) (Laskowski *et al*. 2018). The iPfam database was a database of protein domain-domain and domain-ligand interactions based upon domain annotations from the Pfam domain database, and focussed on interactions directly observed in a structure, rather than mapping ligands to their cognate counterparts (Finn *et al*. 2014). BioLiP2 is a database of protein-ligand interactions specifically focused on identifying biologically relevant interactions through an artefact list of approximately 500 common artefact ligands (Zhang *et al*. 2024).

The PROCOGNATE database mapped domain-cognate ligand interactions to extract the biological relevance of domain-ligand interactions (Bashton, Nobeli and Thornton 2008). It included domain annotations from CATH, SCOP, and Pfam to provide both structural and sequence domain annotations, together with cognate ligand annotations from KEGG. These mappings have been used for evolutionary studies of domain and cofactor origins (Caetano-Anollés, Kim and Caetano-Anollés 2012), to filter structures utilised in stability studies to only those containing cognate ligands (Juritz *et al*. 2012) and as a tool to curate collections of cognate ligands for other databases (Lopez *et al*. 2011). This database is no longer maintained, and in the meantime the PDB has continued to grow, adding to the number of structures without cognate ligand assignments.

Many of PROCOGNATE’s former features exist in other databases, for example, PDBSum contains cognate ligand similarity and contacts between protein residues and the PDB ligand, and BioLiP2 annotates ligands and their receptor binding sites as well as filtering of non-biologically relevant ligands through a curated list of additives. Whilst one or many of the original PROCOGNATE’s features have been replicated in subsequent databases, no complete cognate ligand-domain mapping has been available since the deprecation of PROCOGNATE after version 1.6 - motivating the creation of our new database of cognate ligand mappings.

At all scales, biology can be seen as highly interconnected; for example - microbiomes are complex mixtures of microbes working cooperatively and competitively (Ma *et al*. 2020); at the cellular level, communication between cells is essential to life (Jin *et al*. 2021); and at the subcellular level, proteins rely on networks of interactions between amino acids to form 3D structures necessary to function, and to interact with their substrates (ligands) (Vazquez *et al*. 2003). Therefore, representing the highly connected nature of protein domain-ligand interactions through a graph database may present the opportunity to draw new understandings from these interaction networks - for example, understanding the central cognate ligands in biology and how particular types of domains combine to interact with ligands.

To this end, we present a new analysis pipeline and graph database of cognate ligand-domain interactions, utilising up-to-date methods and a more comprehensive range of compound and reaction databases to explore the ever-expanding set of structures available in the wwPDB and a flexible user interface to easily harness the knowledge of the underlying graph database.

## 2 Implementation and Methods

### 2.1 The ProCogGraph Pipeline

ProCogGraph integrates data from SIFTS (Velankar *et al*. 2013), the Chemical Component Dictionary (Westbrook *et al*. 2015), PDBe-modelserver (Varadi *et al*. 2022) and various reaction and cognate ligand structure databases. A NextFlow (Di Tommaso *et al*. 2017) pipeline is utilised to build the database from a target structure manifest and cognate ligand dataset (Fig. S1). Enzyme structures are downloaded from relevant webservers (step 1a) using PDBe-KB preferred assemblies (accessed from https://ftp.ebi.ac.uk/pub/databases/pdbe-kb/complexes/assemblies_data.csv) and SIFTS EC annotation mapping (accessed from https://ftp.ebi.ac.uk/pub/databases/msd/sifts/csv/pdb_chain_enzyme.csv). Updated MMCIF files and SIFTS XML files are downloaded from the PDBe. Protonated assemblies are obtained from PDBe-modelserver, which enables hydrogen bonds to be determined within structures.

Cognate ligands are aggregated from KEGG, ChEBI, RHEA, GlyTouCan and PubChem (Morgat *et al*. 2015; Hastings *et al*. 2016; Kanehisa *et al*. 2017; Kim *et al*. 2018; Fujita *et al*. 2021) (step 1b), using KEGG reaction IDs to map cognate ligands to EC reactions. SMILES strings are obtained for the various ligands, and these structures are loaded and processed using RDKit v2024.03.2 (Landrum *et al*. 2024). Cognate oligosaccharides are obtained through database cross-referencing between KEGG compound records and the GlyTouCan database GlycoCT sugar encoding. This is converted to a SMILES structure using the Glycan-FormatCoverter and CSDB conversion APIs (Chernyshov and Toukach 2018; Tsuchiya, Yamada and Aoki-Kinoshita 2019). Cognate ligands are annotated with their cofactor status based on their ChEBI cross-references and their role, as assigned by ChEBI ontology. A benefit of the cognate ligand curation approach taken is that cognate ligands are collected for all EC numbers and not just those with a protein chain mapping, allowing for easy integration of new structures to the database in future releases. Bound entities (bound ligands and branched oligosaccharide entities) are identified in the structure from the updated mmCIF file. SMILES representations of bound ligands are obtained by cross-referencing ligand PDB codes to the Chemical Component Dictionary. A similar process to cognate oligosaccharide representations is employed for oligosaccharides in PDB structures. Where available, the WURCS representation of an oligosaccharide is obtained from the `_pdbx_entity_branch_descriptor` field and converted to SMILES as described above.

The Proportion of Atoms Residing in Identical Topology (PARITY) method is used to measure similarity between bound entities and cognate ligands (Tyzack *et al*. 2018). A structural similarity method was chosen for use in the database due to the anticipation that structures being compared for cognate similarity can be reasonably expected to be similar to each other if they are cognate, and using a structural similarity method in these situations allows for more sensitive assessment of the similarity between ligands than chemical fingerprints. The PARITY method provides a fast, easily interpretable similarity between two ligands whilst allowing for small deviations in the maximum common structure between the two, preventing over-penalisation of highly similar structures with small atom differences in the MCS (1 = identical, 0 = no similarity). Various cutoffs for PARITY score have been used previously (Tyzack *et al*. 2018; Špačková *et al*. 2024), and in ProCogGraph, a minimum threshold is determined through analysis of random ligand pairs, to minimise the number of spurious matches. Five sets of 2,000 randomly paired ligands were retrieved, and PARITY similarity calculated, with the 95th percentile score determined for each set. The mean 95th percentile score represents the typical similarity of non-cognate ligand pairs (0.29±0.01). To confidently assign a cognate ligand in ProCogGraph, the minimum required PARITY score is set 10% above this mean 95th percentile at 0.4 PARITY score. An important design decision in ProCogGraph is not to exclude multiple cognate ligands for the same bound entity. The minimum similarity threshold described above removes poor-quality matches; however, where multiple valid matches are available for a ligand, all results are presented to allow users to determine the appropriate “best” cognate ligand.

Contacts are determined using PDBe-arpeggio (Jubb *et al*. 2017) and mapped to protein domains from CATH, SCOP, Pfam, Super-family, Gene3DSA and SCOP2 (Wilson *et al*. 2009; Lees *et al*. 2012; Andreeva *et al*. 2020; Blum *et al*. 2021; Mistry *et al*. 2021; Waman *et al*. 2024) using SIFTS annotations (step 2, PROCESS_CONTACTS). A minimum residue cutoff of 3 filters spurious contacts between residues and PDB ligands. This cutoff ensures that peripheral ligands to the assembly and sparse domain interactions with a ligand are removed, such as the DNA intercalating agent EVP, which binds to DNA in PDB 8J9V, and has a small number of contacts with the protein entities in the structure. Following the assignment of contacts, the domain interaction mode is assigned according to a set of cutoffs determined through evaluation of the distribution of contacts between domains and ligands. Interactions are based on the percentage of overall contacts identified to the ligand, regardless of type. This data is available within the graph as a property of the INTERACTS_WITH relationship. In addition, residue-level interactions between domains to a ligand are stored, providing a summary of the ligand binding interface of a domain.

Previously, PROCOGNATE used a binary interaction mode for protein domains and ligands, assigned as either “shared” or “non-shared” (Bashton, Nobeli and Thornton 2006). Analysis of the distribution of contacts from domains (Fig. S2) shows that ligands are predominantly bound by a single domain, with 2 and 3-domain interactions present at a much lower level and 4+ domain interactions highly infrequent. One or two domain interactions make up >96% of all ligand interactions in ProCogGraph, across all domain database types (in the CATH database, single domain interactions make up 77% of all domain interactions). Gini index was used to investigate the inequality of interaction in multi-domain ligand interactions, which is commonly employed in financial settings to measure income inequality (Hasell 2023), and has been used in bioinformatics to describe gene expression level distribution (O’Hagan *et al*. 2018) and in chemical probe selection (Ursu *et al*. 2020), amongst others. A Gini value of ≥0.4 is commonly accepted to represent relative inequality, and the mean lowest domain contact percentage for interactions exhibiting relative inequality was 6.9% for 2-domain interactions, 7.5% for 3-domain interactions and 4.7% for 4-domain interactions in our analysis. On this basis, we set a “minor” domain interaction as any domain contributing fewer than 10% of contacts to a ligand. Minor interactions were verified by assessing the energetic (un)favourability of domain interaction using Surfaces (Teruel, Borges and Najmanovich 2023). Ten ligands containing “minor” interactions were randomly selected, together with ten ligands containing interactions between 10 and 20% of interactions, to represent the next closest group of domain contacts to the “minor” domains. Domains with 10-20% (low partner) contacts have a significantly (p < 0.001) stronger interaction energy with ligands than minor domains (Fig. S3). Where all except one domain interaction is classified as minor, this single domain is classified as either dominant (≥90%) or major (10-90%). In ProCogGraph, we extend the “shared” definition of PROCOGNATE to increase the granularity of partner interactions. Two additional contact categories are defined to characterise these interactions - major partner (≥50% of all contacts within a ligand interaction) and partner (≥10% -<50% contacts). Table S1 contains definitions for all domain interaction modes utilised in ProCogGraph, and their abundance in ProCogGraph relative to the total number of domain interactions for each domain database. Major partner and partner domains are the second most abundant interactions observed in ProCogGraph, behind exclusive interactions.

Interactions of this type may, for example, be cases of active site formation between domains or interactions where one domain is specialised to hold one moiety of a ligand and another holds a different moiety - see section 3.5 for further exploration of combinatorial domain interactions. ProCogGraph uses Neo4j v5.22.0 community edition as a database management system (DBMS), and NeoDash v2.4.8, a Neo4j plugin, is used for the dashboard interface. To ensure the interoperability of ProCogGraph regardless of the DBMS, ProCogGraph is distributed in flat file format alongside a build script for building the Neo4j version of the database, that can easily be adapted to other DBMS (Fig. S4 gives an overview of the graph schema). The ProCogGraph mappings are also distributed in a flat-file format to allow for custom data exploration and integration with other tools and research. A domain-cognate ligand mapping file and domain-bound entity mapping file are provided for each domain database type. For integration of the cognate ligand similarity data into other tools and analyses, a PARITY similarity mapping file between PDB ligands and cognate ligands is also distributed. For details on running the ProCogGraph pipeline, database and dashboard, see Supplementary Text Section 1.

### 2.2 Analysis Using ProCogGraph

Graph queries are performed using the Cypher language via the Neo4j Python driver and processed using Pandas v2.2.2. Where SMILES strings were returned from a query, these were processed using RDKit to generate Mol objects. Cognate ligand similarity space was explored using ligand MACCS 166 key fingerprints (Kuwahara and Gao 2021), generated via RDKit. These fingerprints were visualised using t-SNE (Maaten and Hinton 2008) to reduce dimensionality to 2D, with projections clustered using k-means clustering and the appropriate cluster number established through Silhouette analysis using scikit-learn v1.5.0 (Pedregosa *et al*. 2011). Similarity of cognate ligands was assessed using Tanimoto similarity of fingerprints at a threshold set based on analysis by Landrum using ligands from ChEMBL (Landrum 2021) demonstrating a Tanimoto threshold of ≥0.575 accounts for 90% of randomly paired molecules (strict threshold herein) and a threshold of ≥0.431 accounts for 70% of random pairs (permissive threshold herein). A promiscuous domain is defined in our analysis as a domain which has non-minor interactions with four or more cognate ligands, a threshold previously described, which avoids matching domains which interact with a cofactor, substrate and product from a single reaction (Bashton, Nobeli and Thornton 2006).

## 3 Results

Fig. 1 depicts the breakdown of structure annotation in ProCogGraph. Many structures are not enzymes/do not contain bound entities, and so are omitted. Overall, 95% of structures that can be annotated are included in the database. Table S2 summarises the domains, bound entities, and cognate ligand mappings in ProCogGraph compared to the last available version of PROCOGNATE (v1.6, March 2009). ProCogGraph significantly increases coverage, with four additional domain data sources added, an 11.2x increase in PDB structures, and a 4.5x increase in bound ligands with cognate ligand mapping.

**Fig. 1.**
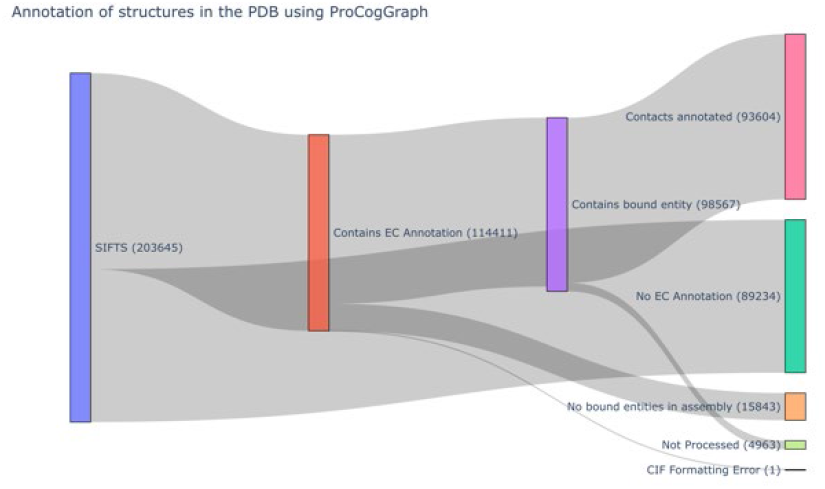
Procession of structures through the ProCogGraph pipeline. At each stage, some structures are lost due to either: no EC annotation, no bound entities or domains in structure, or structure contacts failing to meet criteria.

### 3.1 The ProCogGraph Dashboard

ProCogGraph is also accessible through a dashboard, which enables a rich interrogation of protein domain - cognate ligand mappings for all annotated structures and search functionalities by EC ID, cognate or bound entity, and protein domain. The dashboard allows users to filter matches by similarity score, access specific domain databases e.g. CATH, SCOP2, Pfam, and filter interactions to, for example, only view cognate ligand interactions with a domain which are exclusive. A detailed description of the ProCogGraph dashboard is available in Supplementary Text Section 2 and Fig. S10.

### 3.2 Cognate Ligands in ProCogGraph

At the minimum similarity threshold, 169,693/489,661 bound entities (representing 7,408 distinct structures) have 1+ mappings to cognate ligands, of which 73,590 are perfect matches. In line with previously reported cognate ligand mappings (Bashton, Nobeli and Thornton 2006), more than 50% (41,945) of these matches are to cofactors. To understand the chemical space cognate ligands occupy, assigned cognate ligands were clustered using their chemical fingerprint by k-means clustering (n=14, mean silhouette coefficient 0.51 - see Fig. S5). Some well-distinguished clusters are apparent within the projection, including clusters 4, 5, and 11-13. Inspection of the ligands within these clusters shows that they correspond to: nucleotides/derivatives (cluster 11), sugars (cluster 5), phosphorylated sugars and metabolites (cluster 4), pyrroles (cluster 12), and Coenzyme A/conjugates (cluster 13). Additionally, cluster 9 (silhouette score 0.45) contains many amino acids and their intermediates, such as L-saccharopine, a compound produced in the degradation of lysine.

The cognate ligand chemical space (Fig. 2) reflects the diversity seen in assigned perfect matches: many cognate ligands cluster closely together and seem to cluster based on their core cofactor structures e.g. pyrroles (cluster 12) or Coenzyme A (cluster 13), with the remaining clusters less well defined and representing the overall product/substrate space of cognate ligand binding. The clustering of ligands is used in subsequent sections to relate chemical similarity of cognate ligands to domain binding.

**Fig. 2.**
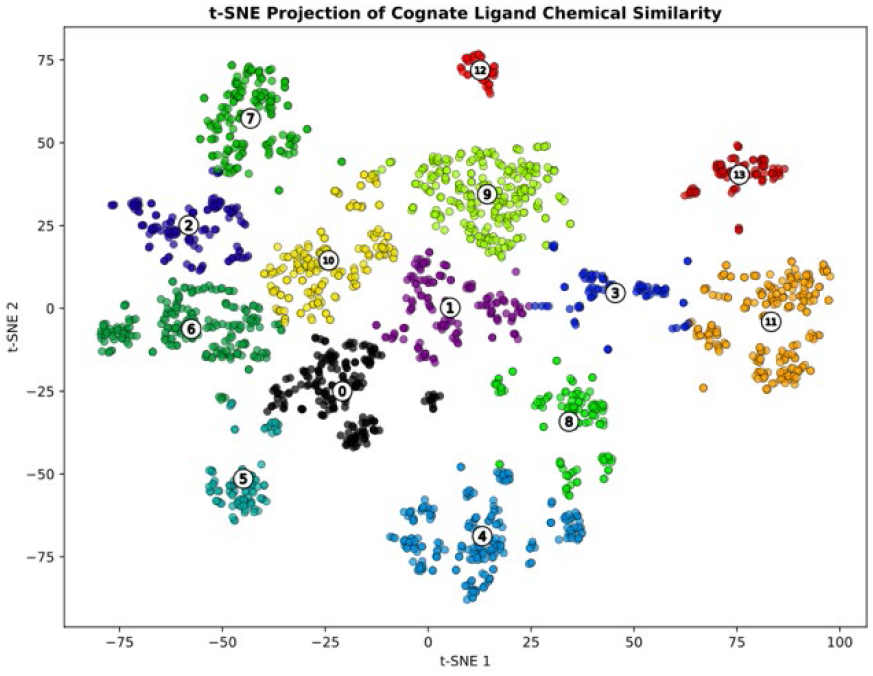
Cognate Ligand Cluster Visualisation using t-SNE. Each point represents a unique cognate ligand, which is coloured according to its assigned cluster using KMeans clustering. Cluster centroids are annotated with the cluster number. Not all clusters within the ordination have clearly similar structure or function - however it can be seen that cluster 4 corresponds to phosphorylated sugars and metabolites, cluster 5 to sugars, cluster 9 to amino acids/intermediates, cluster 11 to nucleotides/derivatives, cluster 12 to pyrroles and cluster 13 to Coenzyme A/conjugates. The remaining clusters are defined as the cognate substrate/product space.

Only 25% of the total number of bound entity descriptors (7,408/29,635) have a mapping to a cognate ligand. Ligands that are never mapped may be genuine non-cognate ligands, map to a cognate ligand not present in any of the cognate ligand databases, or not meet the minimum similarity threshold. Of the 22,086 bound entity descriptors with no cognate ligand match, the top 10 ligands account for 8% of all bound entity occurrences of these descriptors, and include common membrane components and ions used as part of substrate analogues (Supplementary Text Section 3 and Table S3).

### 3.3 Chemical Diversity in Domain-Ligand Interactions

Within ProCogGraph, 460 promiscuous superfamilies are identified. One aspect of cognate ligand interaction left unexplored by simply considering the promiscuity of a domain interaction is the chemical (dis)similarity between the cognate ligand set. To address this, the within and between group chemical diversity of cognate ligands bound by superfamilies was explored using cognate ligand fingerprints. Superfamilies can be explored as either “specialised” - where the superfamily interacts with highly similar ligands only, or “generalised” - where the cognate ligands a superfamily interacts with show a high degree of dissimilarity.

Many superfamilies interact with a small number of cognate ligands - 763/1223 (62.4%) superfamilies interact with three or fewer cognate ligands. In such cases, it may be because of a very specialised function or lack of cognate ligand or bound entity representation in the underlying dataset. Of the 460 promiscuous superfamilies, 15.7% interact with chemically similar (Tanimoto ≥0.575) cognate ligands within ProCogGraph. Fig. S6 shows the cognate ligands bound by the superfamilies with the highest intra-group chemical similarity - these specialised domains interact with distinct clusters within the ligand chemical space, particularly those clusters identified as cofactors e.g. cluster 11/12. Conversely, the most “generalised” superfamilies tend to bind across the substrate ligand space (Fig. S7).

Generally, a promiscuous superfamily is also a generalised superfamily. Table S4 shows the most promiscuous superfamilies within ProCogGraph with good agreement compared to the previously reported promiscuous domains defined by PROCOGNATE (Bashton et al. 2006, Table 4). However, examples of promiscuous superfamilies with specialised ligand binding interaction can also be seen. SF 2.40.110.10 (Butyryl-CoA Dehydrogenase, subunit A, domain 2) interacts with 23 potential cognate ligands which have a high (0.75±0.30 Tanimoto score) similarity (Fig. 2 below) - the majority of which are CoA/derivative ligands (cluster 13), with a small subset of FAD/NAD/derivative ligands (cluster 11).

### 3.4 Combinatorial Domain Interactions in ProCogGraph

Domain contexts have been previously explored manually in a select set of circumstances (Bashton, Nobeli and Thornton 2006; Bashton and Chothia 2007), and have been shown to influence the interaction of domains with different cognate ligands. This was investigated by identifying the cognate ligands bound in a “non-shared” manner by a domain and the collection of other domains present in the protein. A combinatorial domain interaction is defined here as the collection of domains (more than 1 domain), at homologous superfamily level for CATH domains, which interact with a given PDB ligand in a non-minor manner, and for which the PDB ligand has a cognate ligand mapping with similarity score ≥0.4.

A total of 1,003 unique domain combinations are observed when using CATH domain data. There are various potential ways in which a combinatorial domain interaction may impact ligand interaction - it may be transient/have no impact on the observed cognate ligands bound in the interaction compared to the individual domains - potentially functioning to bring together a substrate and cofactor or two substrates; it may enable a different or expanded or more specific range of interactions to occur; or improve the interaction of a protein to a ligand also commonly interacted with by one of the partner domains in an exclusive manner.

A duplicate major partner-partner interaction is observed with the superfamily “2,3-Dihydroxybiphenyl 1,2-Dioxygenase, domain 1” (CATH 3.10.180.10). In some structures, this domain interacts exclusively with its bound ligand through a beta-barrel structure from a single domain (Serre et al. 1999) - see Fig. 4A. The duplicate partner interaction observed between two of these domains results in a structure where ligand binding is at the dimer interface, and the two domains each provide beta sheets to the beta barrel-like coordination of the ligand (Fig. 4B).

**Fig. 3.**
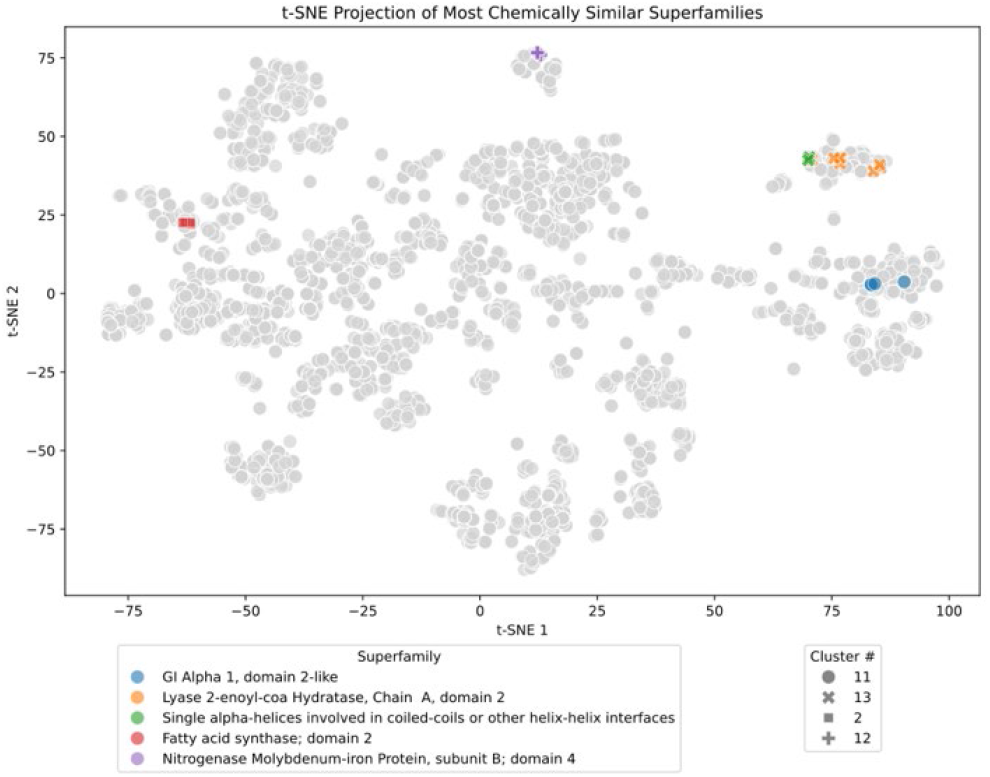
Specialised cognate ligand interactions. t-SNE visualisation of cognate ligands from specialised SF 2.40.110.10 (Butyryl-CoA Dehydrogenase, subunit A, domain 2), coloured, compared to all unique ProCogGraph cognate ligands (grey). Ligands are clustered based on their chemical similarity and coloured according to cluster number. Cluster 13 (orange) consists primarily of CoA/derivative ligands, while Cluster 11 (blue) contains FAD/NAD/derivative ligands with remaining cognate ligands in clusters 0 (green) and 6 (red). The high Tanimoto score (0.75±0.30) indicates a high similarity among the ligands within each cluster.

**Fig. 4.**
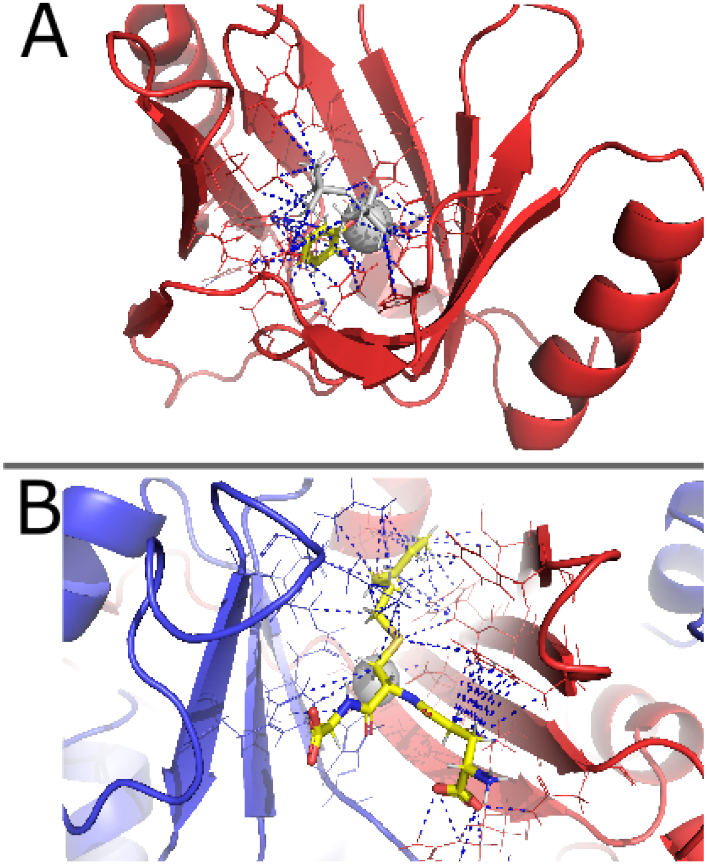
Exclusive and partner interactions of domain 2,3-Dihydroxybiphenyl 1,2-Dioxygenase, domain 1 (CATH Code 3.10.180.10). Blue dashed lines indicate contacts identified by PDBe-arpeggio. A: Exclusive interaction with catechol (stick representation with carbon atoms coloured yellow, cognate = 3-chlorocatechol, 0.89 PARITY score) within the beta-barrel of the domain (PDB 1KND). Zinc ion and tertiary-butyl alcohol ligands are coloured grey. B: Partner interaction of two instances of the domain (PDB 1BH5), coming together to form the enzyme active site for PDB ligand s-hexylglutathione (stick representation with carbon atoms coloured yellow, cognate = (R)-S-Lactoylglutathione, 0.82 PARITY score). Zinc ion highlighted in grey as a space filling representation.

Here, the shared modality of binding enables a larger and more complicated binding pocket at the domain dimer interface, which is able to bind to the larger and more chemically complex glutathione, a tripeptide, through the interface of beta sheets from two domains versus the smaller aromatic diol which is bound in the core of a barrel-like structure in exclusive interactions of the domain superfamily. Fig. S8 shows the topological secondary structure organisation of domains in both 1KND and 1BH5. A possible evolutionary mechanism for this difference is a domain duplication from an original single domain (like that in 1KND) and subsequent divergence modifying the two domains so that they form a functional dimer interface when paired, such as that seen in 1BH5.

Exclusive interactions are observed to aromatic diol cognate ligands such as Naphthalene-1,2-diol and Biphenyl-2,3-diol - to be expected due to the EC annotation of these structures as EC 1.13.11.- (involved in the cleavage of aromatic compounds). Conversely, partner interactions are observed to glutathione and glutamate, in structures annotated as EC 4.4.1.- (carbon-sulfur lyases). Here, glutathione, a tripeptide, is a more complex molecule than the simple aromatic structure bound exclusively and includes multiple functional groups, which may necessitate the larger binding pocket afforded by a multi-domain interaction mode.

A partner interaction between superfamily “Butyryl-CoA Dehydrogenase, subunit A, domain 3” (1.20.140.10) and superfamily “Butyryl-CoA Dehydrogenase, subunit A, domain 2” (2.40.110.10) demonstrates how two domains can act in combination to bind a new cognate ligand. In isolation, these domains interact with ligands such as propanoyl-CoA in a prolyl oxidase structure (PDB 6CY8, EC 1.3.8.1), and 4-hydroxyphenylacetate substrate in 4-Hydroxyphenylacetate 3-hydroxylase (EC 1.14.14.9) structures (CATH 1.20.140.10 PDB 2JBT, CATH 2.40.110.10 PDB 2YYM). In combination, these domains interact with coenzyme FAD, creating a binding pocket, which is used to activate O2 for the incorporation of oxygen into the substrate (Paul et al. 2021). This demonstrates how combinatorial domain interactions can be essential to enzyme function. All three domains are involved in the interaction, but bind distinct components of the FAD molecule. Fig. S9 shows two examples of this coenzyme binding pocket, in a p-hydroxyphenylacetate hydroxylase (Fig. S9A, PDB 2JBS) and in a butryl-CoA dehydrogenase (Fig. S9B, PDB 1BUC). In PDB 1BUC, the adenine moiety of the FAD molecule interacts with residues from superfamily 1.20.140.10 domains in two chains. In contrast, most interactions with the isoalloxazine ring are from a single superfamily 2.40.110.10 domain. Thus, in this instance, the different components of the domain combination each bind a different chemical moiety of FAD.

### 3.5 EC ID completion via cognate ligand detection

ProCogGraph expands cognate ligand comparisons to include all EC IDs for a partial EC annotation e.g. 2.1.1.- (methyltransferases). Using the likely cognate ligand bound by a domain, a specific EC number can be assigned to these structures. 24,041 protein chains across 15,587 PDB structures contain a partial EC annotation, of which 7,923 chains (5,545 PDB structures) have a domain ligand interaction mapping that can be traced to a cognate ligand using our methodology (Figure 5A).

**Fig. 5.**
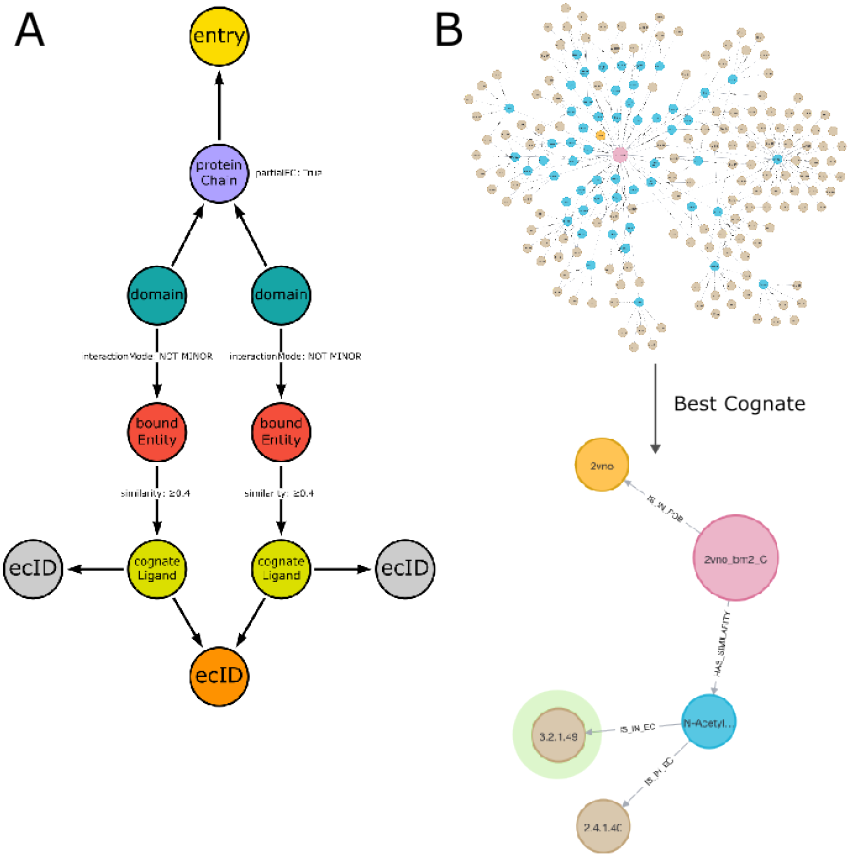
Cognate ligand mapping to determine exact EC IDs. A: The graph schema for matching EC IDs with cognate ligands. For protein chains with partial ECs, non-minor domain interactions to bound entities are traced to cognate ligands with similarity above 0.4. An annotation is made if all cognate ligands share the same EC when multiple bound entities are present. B: For PDB 2VNO, from partial EC 3.2.1, 54 cognate ligands are matched. The highest similarity ligand corresponds to EC 3.2.1.49.

With these mappings, specific EC IDs can be assigned to protein chains by exploring the similarity of all cognate ligands in the undefined EC range to those bound in the PDB structure, improving the annotation of structures in the PDB and allowing greater insight into the functional roles of structures. In ProCogGraph, 1,916 protein chains can be mapped through cognate ligands to specific EC numbers. Occasionally, SIFTS assigns a specific and partial EC number to a protein chain - 604 protein chains with partial EC annotation also contain a specific EC, and so are excluded. This leaves 1,312 protein chains in which the specific enzymatic function of the chain can be assigned through cognate ligand and domain interaction matching. 23.9% (314) of these chains were mapped to the specific EC ID through perfect bound entity-cognate ligand matches, i.e. the bound ligand was the cognate ligand. A full breakdown of the mean similarity score of ligands resulting in EC matches can be seen in Fig. S10.

As an example of the potential of this annotation method, in the PDB structure 2VNO, the assigned EC number via SIFTS was 3.2.1.-, corresponding to Glycosidases, of which there are over 100 EC IDs. The similarity of the bound ligands to cognate ligands from all potential EC IDs is compared, matching 54 cognate ligands across 150 EC IDs. For each bound entity, a “best cognate” property is assigned to the cognate ligand(s) which have the highest similarity score - only 1 cognate ligand has the highest similarity - resulting in an assignment of EC 3.2.1.49, Cleavage of non-reducing alpha-(1->3)-N-acetylgalactosamine residues from human blood group A and AB mucin glycoproteins (Figure 5B).

Often, these EC matches are evident in publications - for example, in PDB 2VNO the specificity of the structure for blood group antigens is discussed. However, using a cognate ligand assignment approach allows systematic annotation of EC IDs. Whilst assigned EC IDs should always be manually verified to ensure accurate mapping, such a method enables large-scale completion of EC identifiers where this cognate ligand mapping is available. This approach works well to assign granular EC IDs to ligands based on observed mappings to cognate ligand substrates/products of reactions and complements other tools for annotation, such as the structure-only based DeepFRI (Gligorijević et al. 2021), enabling the highest-level EC ID annotation where structure alone cannot differentiate between fourth level substrate specific identifiers.

## 4 Discussion

Here, we describe a new cognate ligand-domain interaction database, ProCogGraph, which builds upon its ancestor, PROCOGNATE, by extending coverage 11.22 fold across an expanded set of databases, including cognate ligands from several different databases, incorporating matches to sugars not previously covered, and employing a new data-driven methodology for defining domain-ligand interaction modes and cognate similarity thresholds. Alongside this, a highly customisable dashboard interface is provided, through which users can tailor data to suit their needs, searching for specific structures, EC numbers, cognate ligands or PDB ligands, together with a number of filters for similarity strictness and interaction modes.

We highlight new approaches to considering cognate ligand mapping, exploring the chemical space occupied by cognate ligands, and introducing the idea of specialised domains binding a chemically narrow range of ligands and generalised domains binding a chemically broad range of ligands. We show many of the specialised domains are involved in binding to cofactors, whereas generalised domains are more frequently involved in binding to the substrate/product chemical space. In addition, the combinatorial interaction of domains with ligands and the impact this has on the type of ligand bound is explored. Different mechanisms for combinatorial interaction are highlighted - combinatorial interactions may create binding pockets for new ligands not otherwise seen with exclusive domain interactions or domains may be duplicated, resulting in increased flexibility and altered ligand binding. The knowledge highlighted here could be of use in protein engineering to enable design of proteins able to bind to target ligands flexibly.

We explore the potential reasons for bound entities not matching to cognate ligands and find that many of the most frequently unmatched ligands are common membrane components or compounds used alongside biologically relevant ligands during crystallisation, e.g., beryllium trifluoride with ADP. The most commonly unmatched ligand is Chlorophyll A due to its lack of presence as a cofactor in reaction schemes.

Overall, the cognate ligand-domain mapping ProCogGraph provides is important for continuing to evaluate the core ideas behind cognate ligand domain interactions. It also provides a new interface, metrics, and concepts such as domain generalisation and specialisation, based upon the chemical similarity of cognate ligands, through which to evaluate domains.

## 5 Data and Availability

ProCogGraph is freely available and released under an MIT licence. ProCogGraph source code is available at https://github.com/bash-ton-lab/ProCogGraph and is deposited in Zenodo (https://doi.org/10.5281/zenodo.13249841). Data is archived in Zenodo (https://zenodo.org/doi/10.5281/zenodo.13165851).

## Acknowledgements

The authors thank Dr Layla van Ellen for designing the ProCogGraph logo.

## Funding

This work was supported by Research England’s Expanding Excellence in England (E3) Fund.

### Conflict of Interest

none declared.

---

**Supplementary Table 1:**
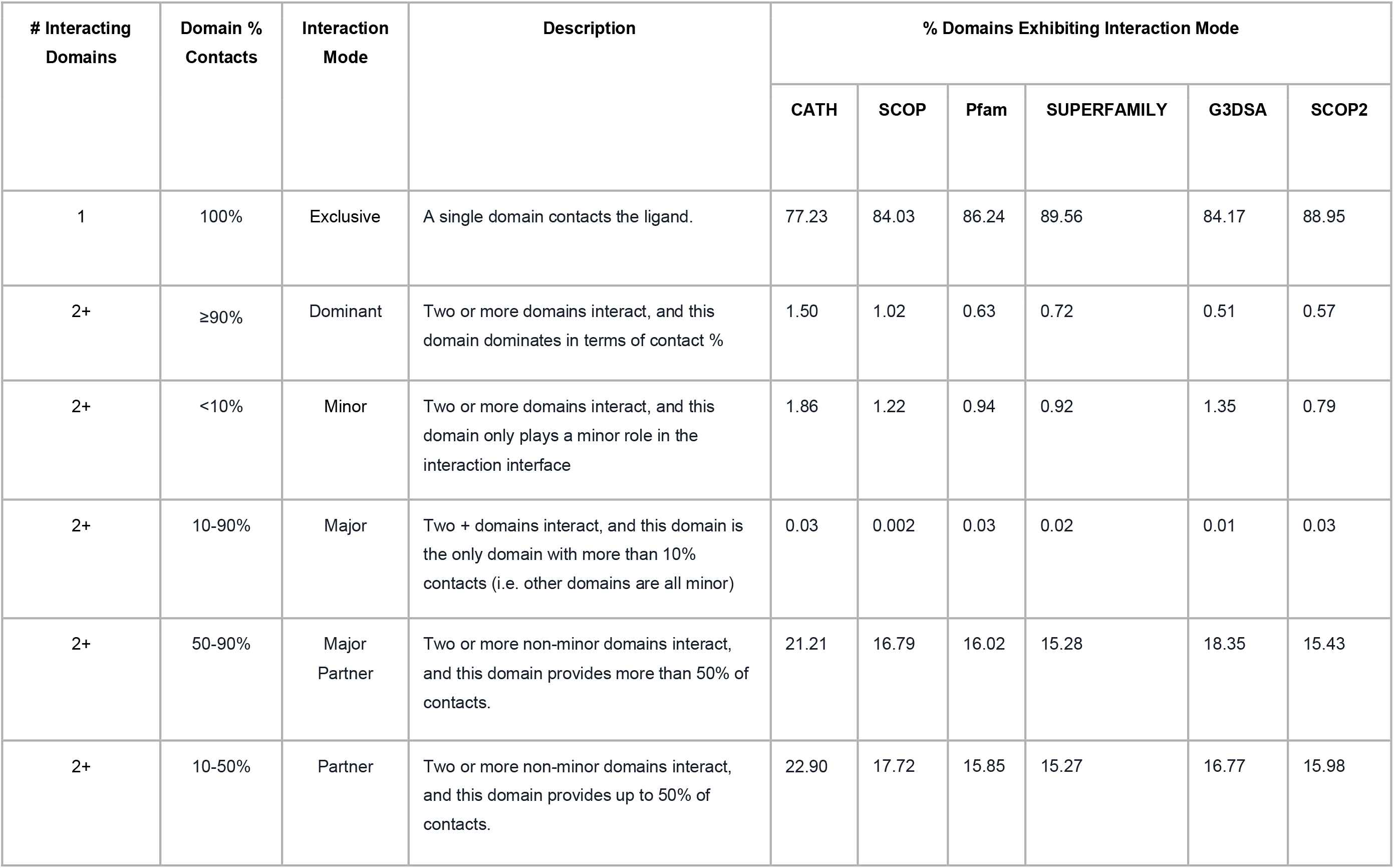
Domain interaction modes in ProCogGraph. One or more domains may interact with a ligand, and depending on the percentage of contacts a domain provides to the interaction, a different interaction mode is assigned. Note one domain may interact with multiple ligands with different interaction modes.

**Supplementary Table 2:** Comparison of PROCOGNATE v1.6 (PCN) and ProCogGraph v1.0 (PCG) coverage for various domain databases. For CATH (v4.2.0) and Gene3D (via InterPro v96.0) databases group level is Homologous Superfamily, for SCOP (v1.75), SCOP2 (v2022-06-29) and Superfamily (via InterPro v96.0), the group level is Superfamily and for Pfam (via InterPro 96.0) the group level is Family. Note: SCOP (v1.75) was last updated in June 2009, resulting in lower coverage.

|  | CATH |  | SCOP 1.75 |  | Pfam |  | SUPERFAMILY |  | Gene3D |  | SCOP2 |  |
| --- | --- | --- | --- | --- | --- | --- | --- | --- | --- | --- | --- | --- |
|  | PCN | PCG | PCN | PCG | PCN | PCG | PCN | PCG | PCN | PCG | PCN | PCG |
| **PDB Entries** | 5,553 | 61,473 | 6,275 | 17,166 | 7,895 | 89,805 | n/a | 74,473 | n/a | 60,500 | n/a | 64,945 |
| **Num Groups** | 427 | 1,907 | 347 | 762 | 7,80 | 3,960 | n/a | 874 | n/a | 1,148 | n/a | 1,054 |
| **EC numbers (only absolutely defined)** | 750 | 2,831 | 873 | 1,694 | 996 | 3,282 | n/a | 2,775 | n/a | 2,435 | n/a | 2,292 |
| **EC Numbers (including expanded partial)** | n/a | 6,652 | n/a | 6,313 | n/a | 6,689 | n/a | 6,651 | n/a | 6,648 | n/a | 6,606 |
| **Ligands In Database** | n/a | 297,547 | n/a | 76,446 | n/a | 433,442 | n/a | 359,828 | n/a | 293,741 | n/a | 309,522 |
| **Ligands with Mapping (0.4 cutoff)** | 24,337 | 104,989 | 26,948 | 28,823 | 32,450 | 151,529 | n/a | 125,909 | n/a | 102,807 | n/a | 101,297 |
| **Number of cognate ligands mapped** | n/a | 2296 | n/a | 1,189 | n/a | 2710 | n/a | 2,340 | n/a | 2,098 | n/a | 2,114 |

**Supplementary Table 3:**
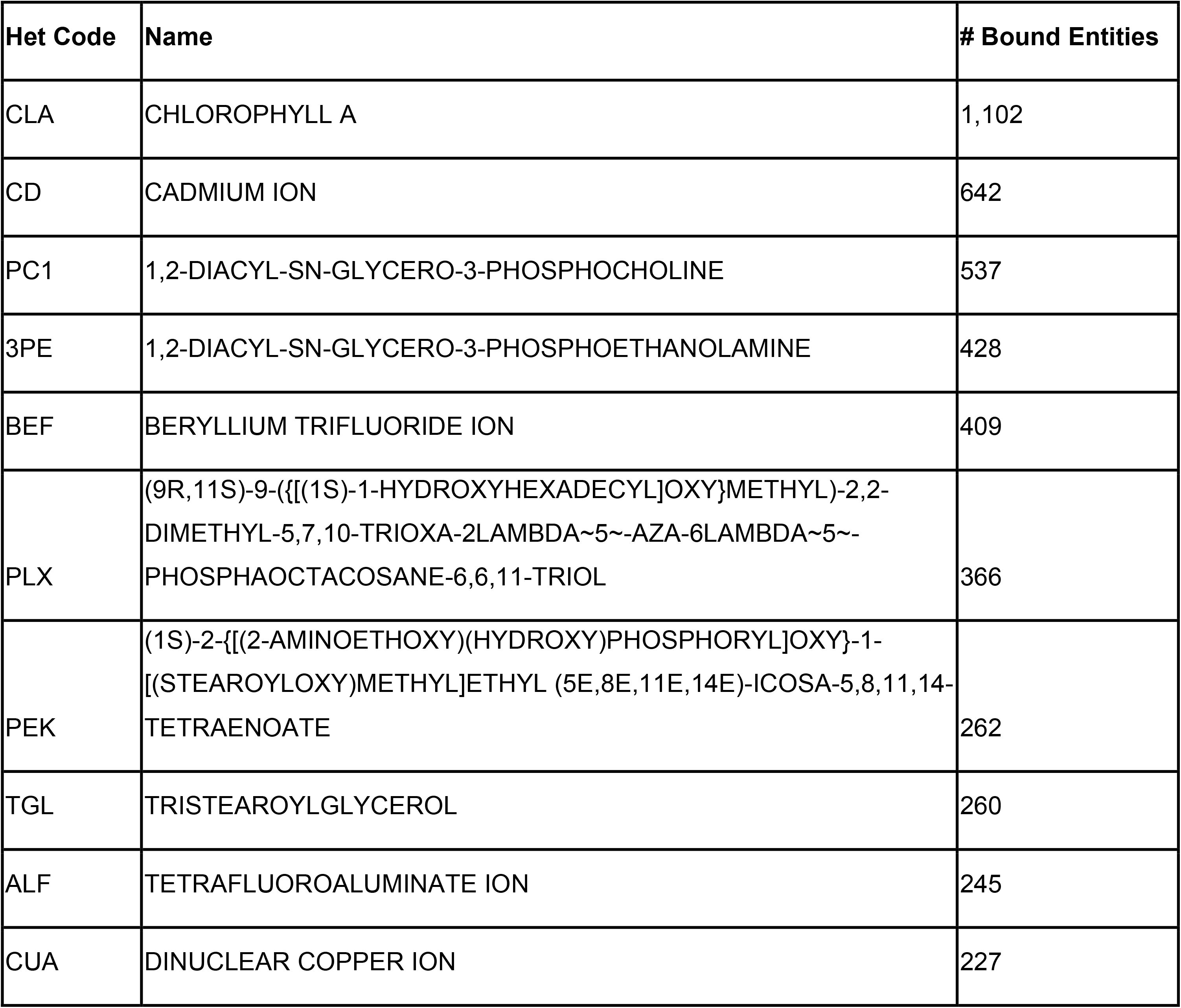
Top 10 PDB Ligands with no cognate ligand mapping in ProCogGraph.

**Supplementary Table 4:** Top 20 most promiscuous domains as defined by number of different cognate ligands a superfamily interacts with.

| **CATH Homologous Superfamily** | **# Cognate Ligands** | **Mean Similarity** | **Similarity St. Dev.** |
| --- | --- | --- | --- |
| NAD(P)-binding Rossmann-like Domain | 338 | 0.29 | 0.19 |
| Aldolase class I | 186 | 0.31 | 0.20 |
| NADP-dependent oxidoreductase domain | 157 | 0.30 | 0.19 |
| alpha/beta hydrolase | 153 | 0.25 | 0.16 |
| Cytochrome P450 | 139 | 0.27 | 0.19 |
| FAD/NAD(P)-binding domain | 128 | 0.26 | 0.18 |
| P-loop containing nucleotide triphosphate hydrolases | 116 | 0.33 | 0.25 |
| Vaccinia Virus protein VP39 | 108 | 0.28 | 0.18 |
| Type I PLP-dependent aspartate aminotransferase-like (Major domain) | 100 | 0.35 | 0.17 |
| **Glycosidases** | 91 | 0.48 | 0.23 |
| 3.20.20.140:Metal-dependent hydrolases | 84 | 0.28 | 0.19 |
| 3.90.1150.10:Aspartate Aminotransferase, domain 1 | 72 | 0.34 | 0.18 |
| 3.90.180.10:Medium-chain alcohol dehydrogenases, catalytic domain | 69 | 0.24 | 0.17 |
| 3.30.360.10:Dihydrodipicolinate Reductase; domain 2 | 68 | 0.35 | 0.19 |
| **3.90.550.10:Spore Coat Polysaccharide Biosynthesis Protein SpsA; Chain A** | 65 | 0.55 | 0.22 |
| **3.90.79.10:Nucleoside Triphosphate Pyrophosphohydrolase** | 57 | 0.51 | 0.34 |
| 3.40.50.620:HUPs | 56 | 0.38 | 0.25 |
| 3.30.420.10:Ribonuclease H-like superfamily/Ribonuclease H | 56 | 0.25 | 0.15 |
| 3.40.630.30:Aminopeptidase | 53 | 0.25 | 0.19 |
| 3.30.70.270:Alpha-Beta Plaits | 51 | 0.30 | 0.20 |

**Supplementary Figure 1:**
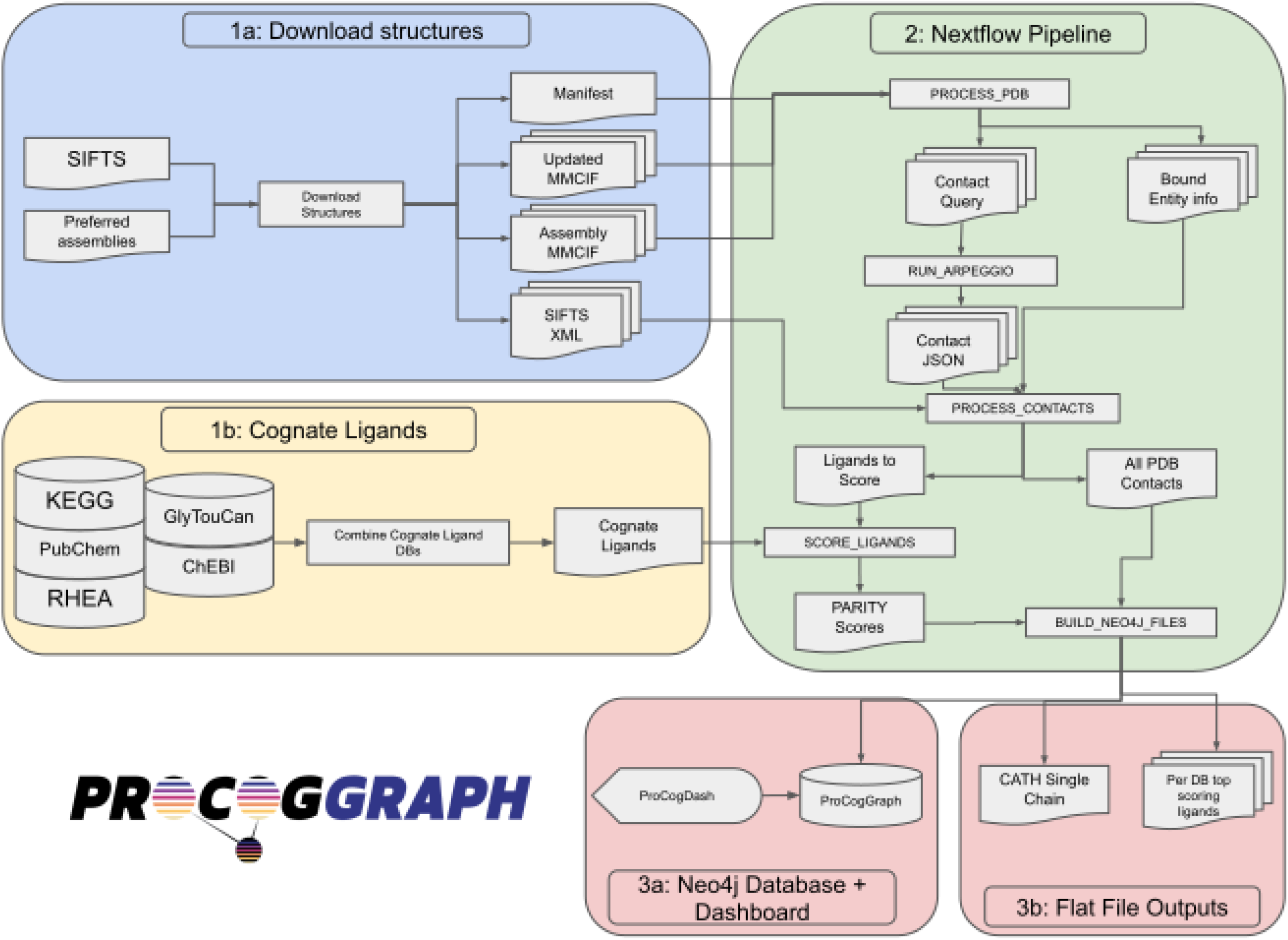
The ProCogGraph Pipeline. 1a) Updated and protonated MMCIF files and SIFTS XML annotations are downloaded from EBI servers (PDBe and modelserver). 2) The Nextflow pipeline is executed, processing each PDB structure to determine bound ligands in the assembly and generate query files for Arpeggio. The generated ligand contacts are then parsed into domain-specific ranges, and interaction modes between domains and ligands are determined. 3) Finally, Neo4j format files and flat file outputs are generated for the ProCogGraph database.

**Supplementary Figure 2:**
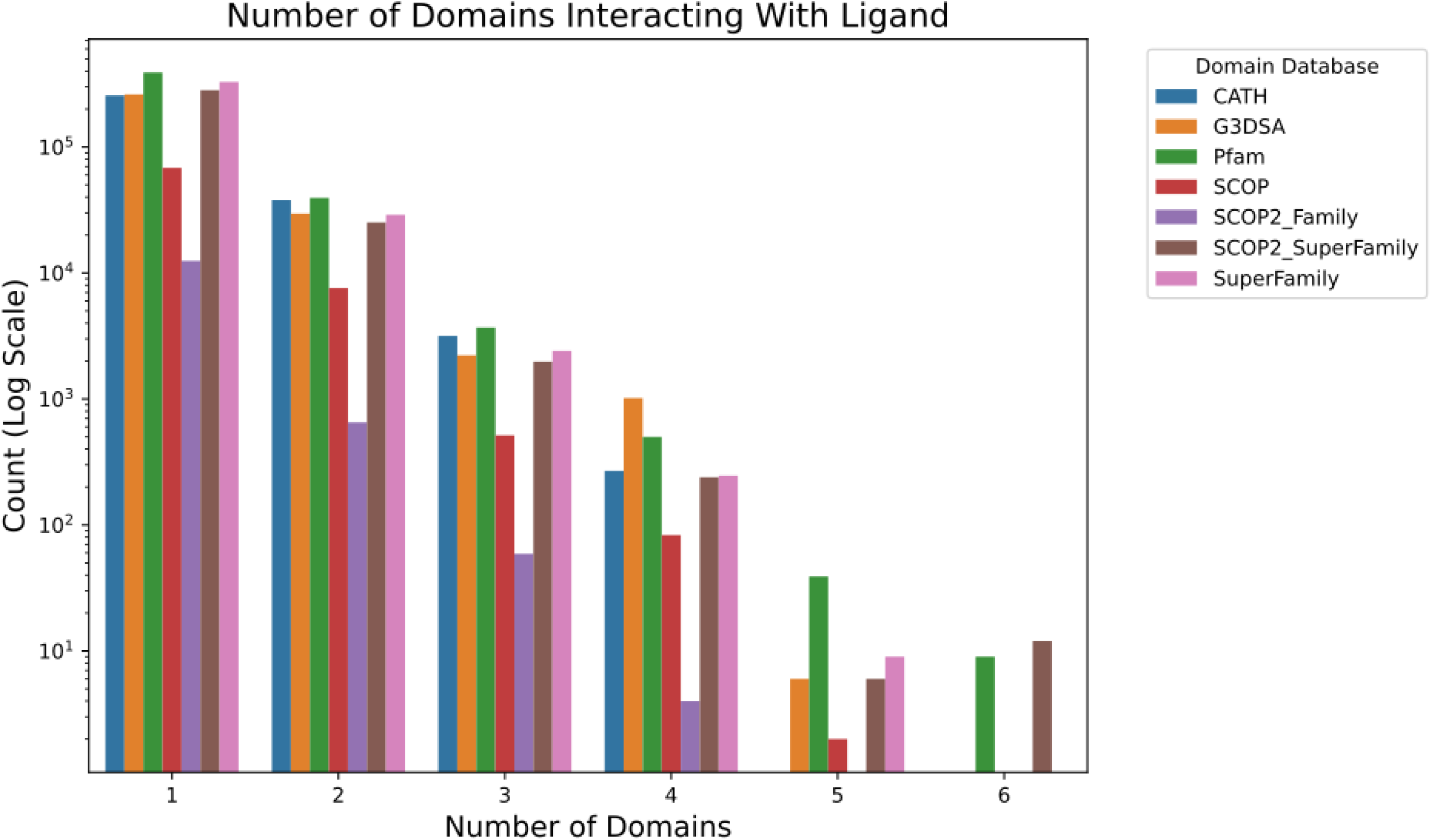
Breakdown of number of domains interacting with bound entities in ProCogGraph, for each domain database. Most interactions to a bound entity involve one or two domains, with 3+ multi-domain interactions less frequent. Five and six domain interactions are observed in a few, but not all domain databases.

**Supplementary Figure 3:**
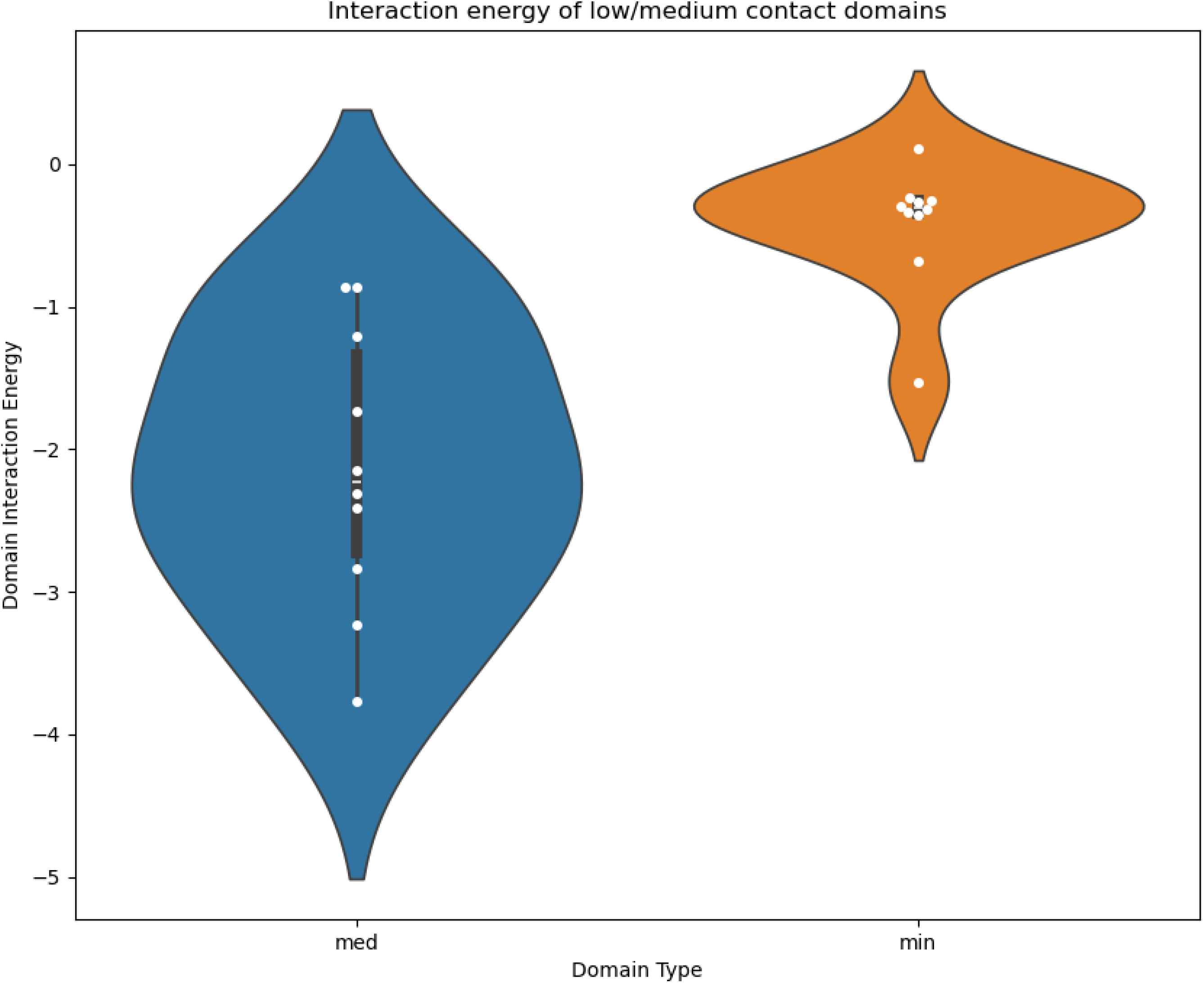
Mean interaction favorability of minor and medium contact domains to their bound ligands.

**Supplementary Figure 4:**
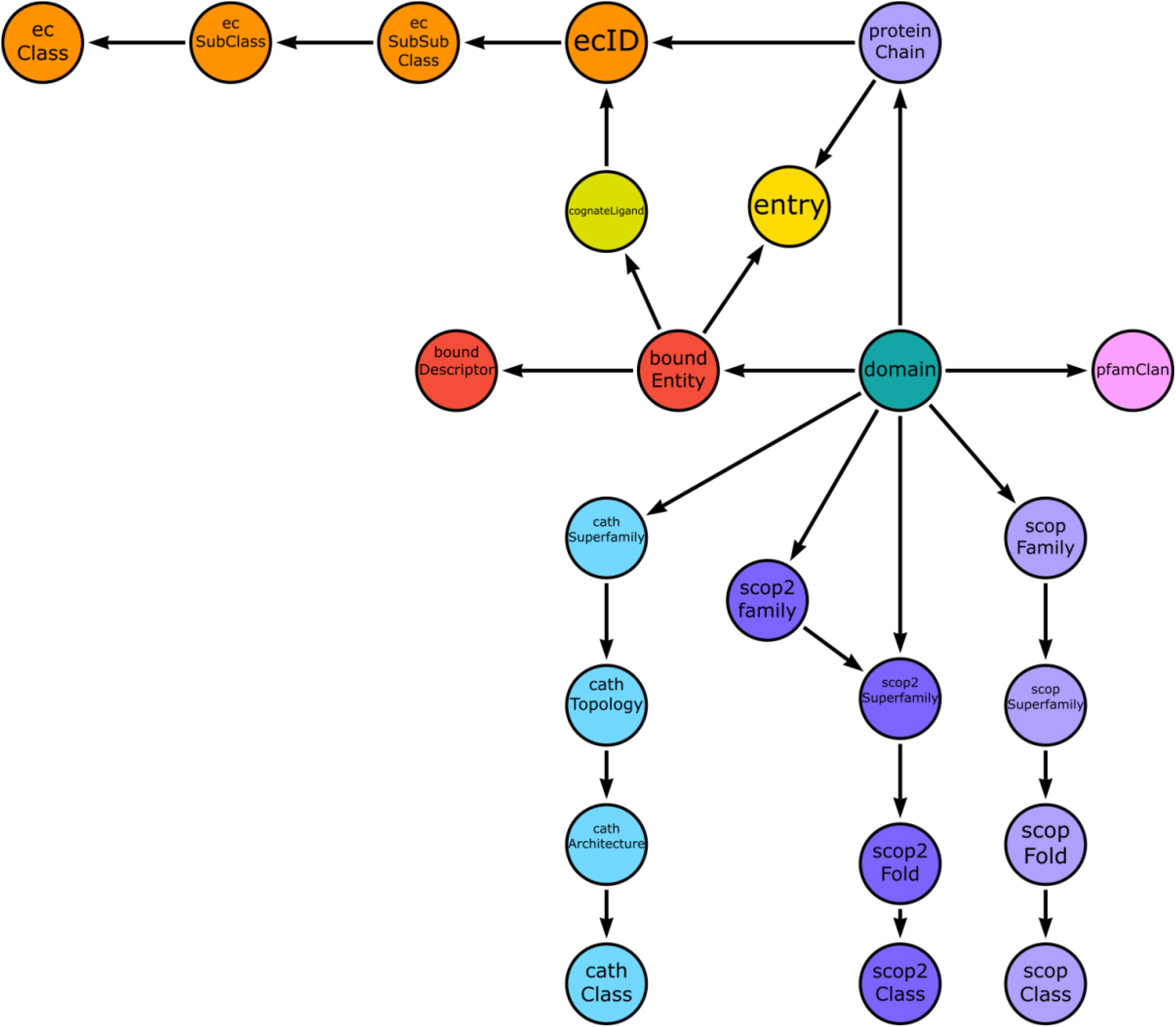
The ProCogGraph schema. The schema can be split at the highest level into four main node types: Enzyme Class, Cognate Ligand, Bound Entity and Domain. Further granularity is then available for the relationships of these nodes, which annotates hierarchical relationships in, for example, the Enzyme Classification system, and also the interrelationship between domains, enzyme classification and cognate ligand enzyme classification.

**Supplementary Figure 5:**
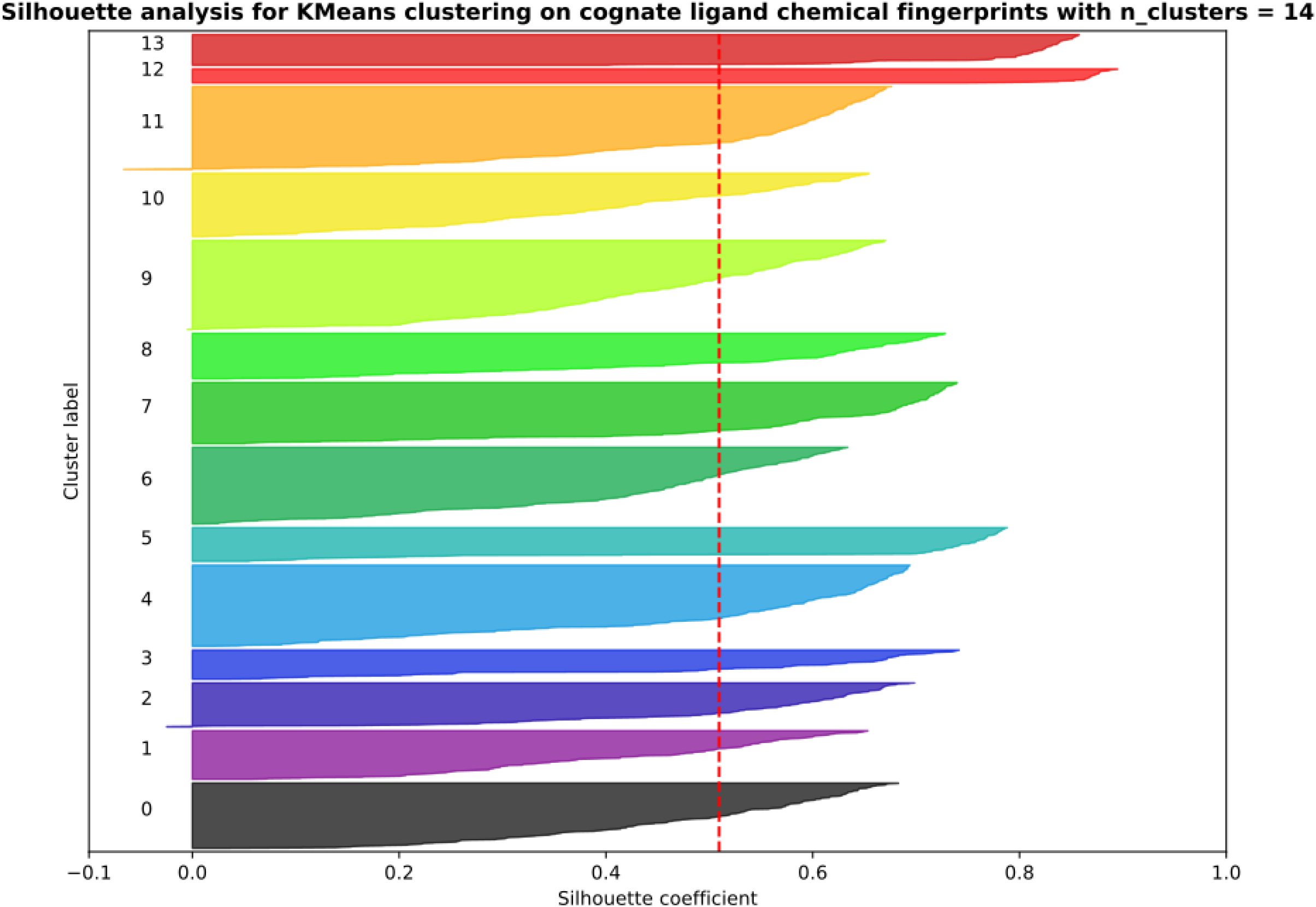
Silhouette analysis of KMeans Clustering on Chemical Similarity of Cognate Ligands, showing the silhouette coefficient for each cluster with n=14 clusters. Red dashed line represents the average silhouette score (0.51) across all clusters.

**Supplementary Figure 6:**
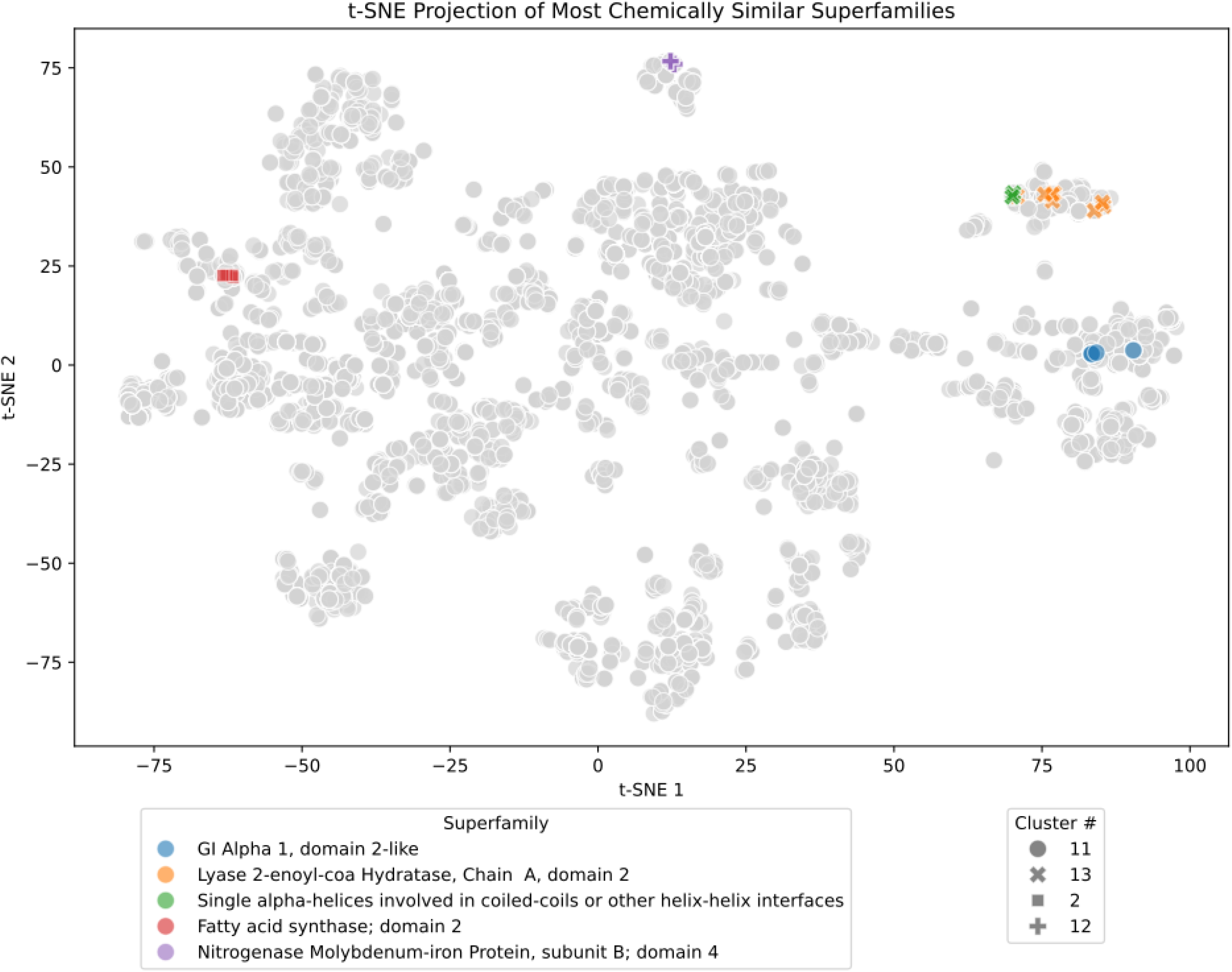
t-SNE visualisation of the five most “specialised” superfamilies with the highest intra-superfamily chemical similarity. Cognate ligands are coloured according to the superfamily they belong to against a backdrop of all unique cognate ligands in the mapping. Ligands from specialised superfamilies are assigned distinct shapes according to the cluster they were assigned to.

**Supplementary Figure 7:**
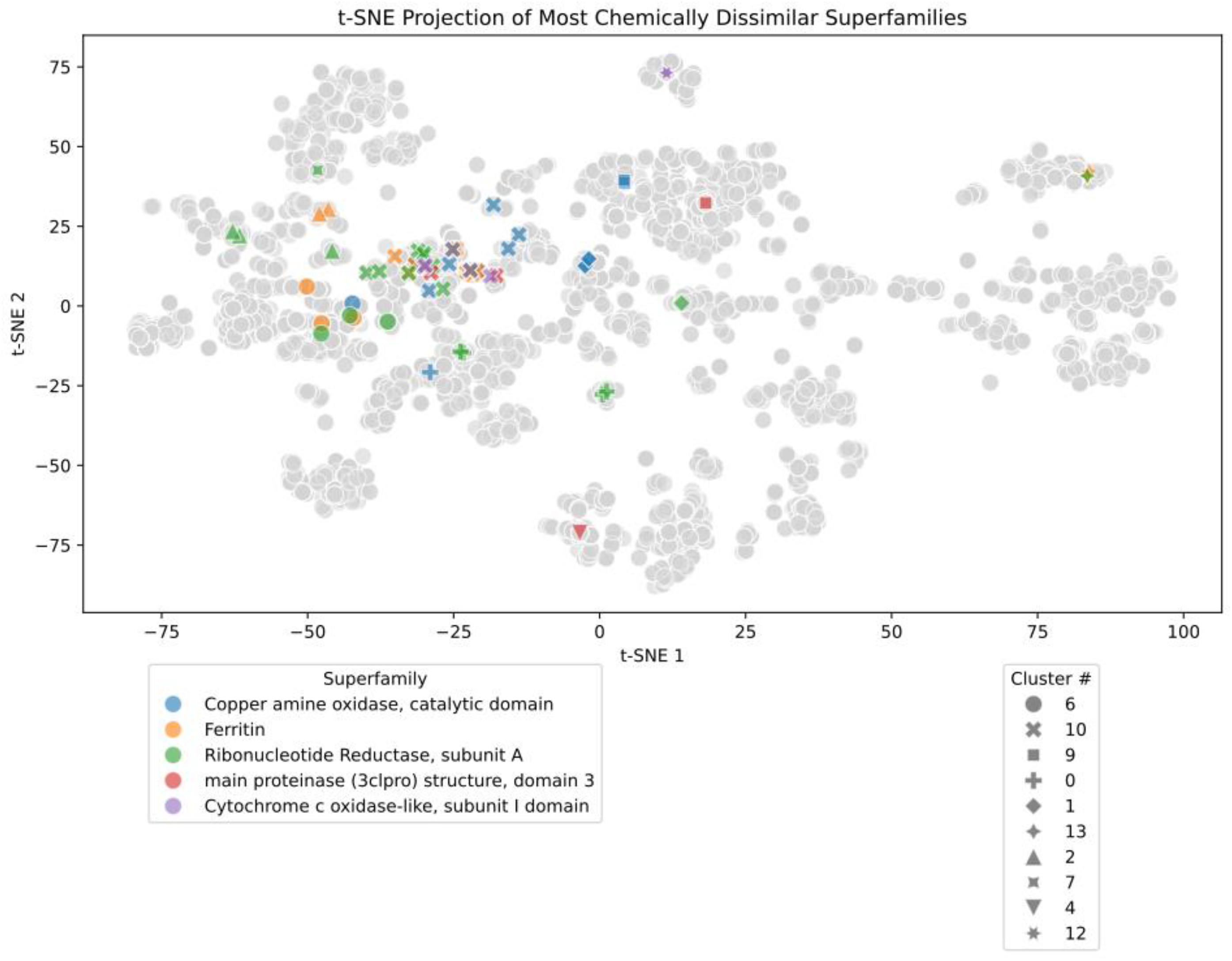
t-SNE visualisation of the five most “generalised” superfamilies with the highest intra-superfamily chemical dissimilarity. Cognate ligands are coloured according to the superfamily they belong to against a backdrop of all unique cognate ligands in the mapping. Ligands from generalised superfamilies are assigned distinct shapes according to the cluster they were assigned to.

**Supplementary Figure 8:**
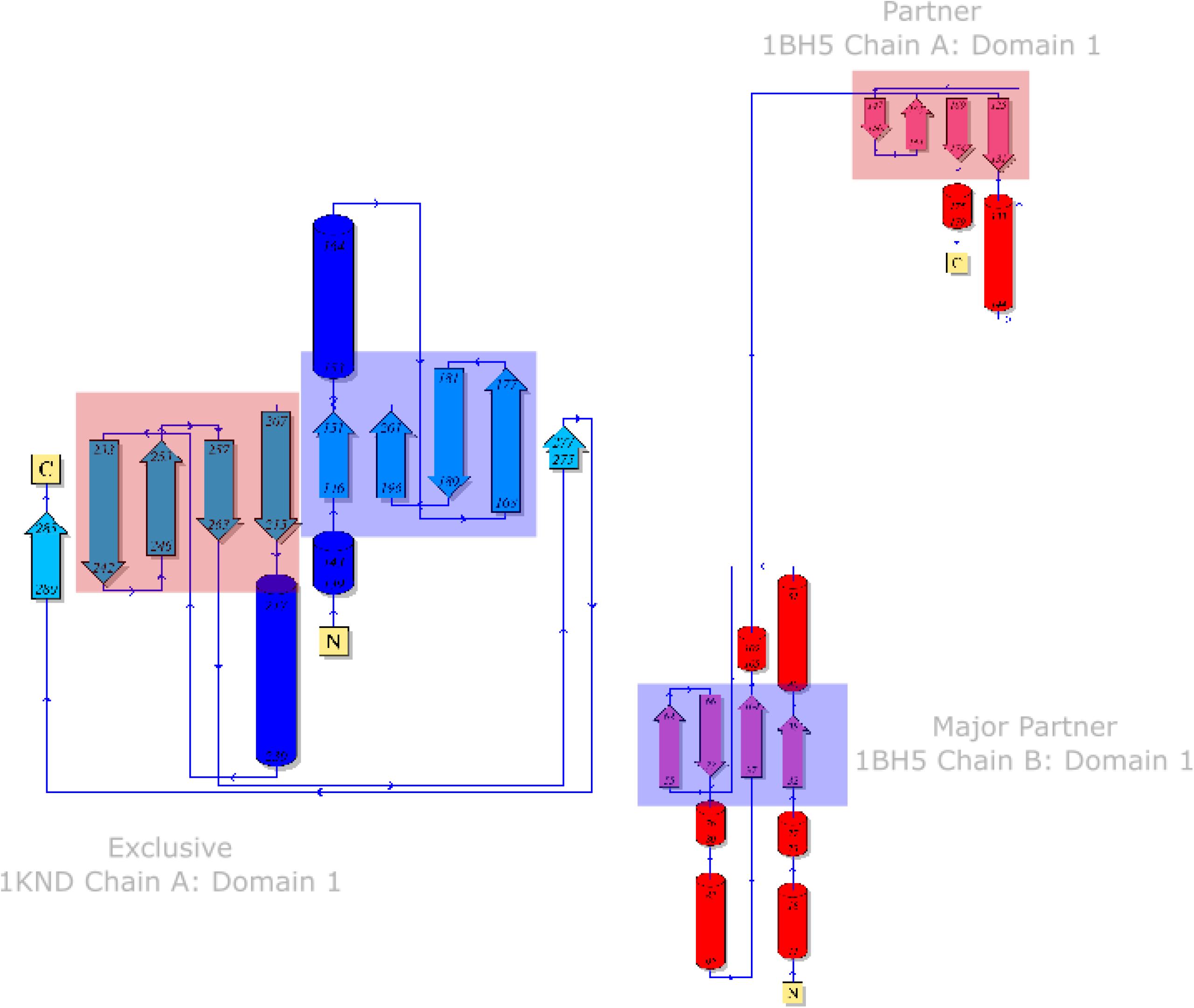
Domain organisation in exclusive and partner binding domains of superfamily 3.10.180.10. Left: Domain organisation in structure 1KND domain 1, which binds it’s ligand exclusively. Right: Domain organisation in structure 1BH5 domain 1, which binds it’s ligand in a partner interaction between two instances of the domain, in chain A and chain B. Blue highlighted beta strands in 1KND domain correspond to beta strands involved in interaction from chain B in 1BH5. Beta strands highlighted in red in 1KND correspond to beta strands involved in interaction from chain A in 1BH5.

**Supplementary Figure 9:**
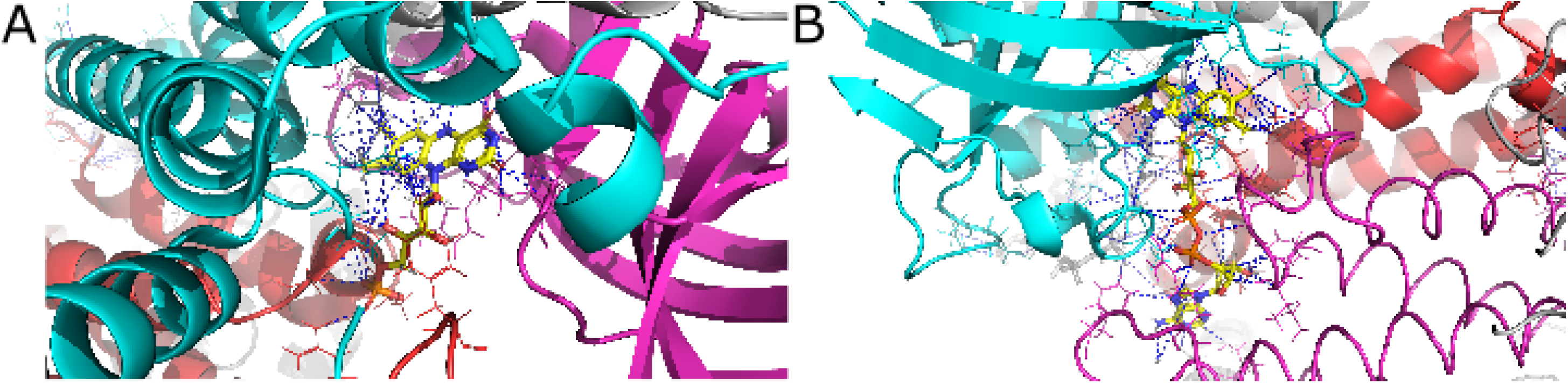
FAD/FMN binding pocket formed by three domains. Interatomic contacts identified between residues are shown with blue dashed lines, residues involved in interaction are shown as wireframe molecules and bound ligands are shown in stick representations with carbons coloured yellow. A: FMN binding pocket within a p-hydroxyphenylacetate hydroxylase (PDB 2JBS). Red: Butyryl-CoA Dehydrogenase, subunit A, domain 3 (CATH 1.20.140.10) domain, chain A. Magenta: Domain Butyryl-CoA Dehydrogenase, subunit A, domain 2 (CATH 2.40.110.10), chain D. Cyan: Butyryl-CoA Dehydrogenase, subunit A, domain 3 (CATH 1.20.140.10) domain. Ligand Flavin Mononucleotide matches to cognate ligand FAD, similarity = 0.59. B: FAD binding pocket within a butryl-CoA dehydrogenase (PDB 1BUC). Magenta: Domain Butyryl-CoA Dehydrogenase, subunit A, domain 3 (CATH 1.20.140.10), Cyan: Domain Butyryl-CoA Dehydrogenase, subunit A, domain 2 (CATH 2.40.110.10), Red: Domain Butyryl-CoA Dehydrogenase, subunit A, domain 3 (CATH 1.20.140.10). Ligand FAD matches to cognate ligand FAD with perfect similarity.

**Supplementary Figure 10:**
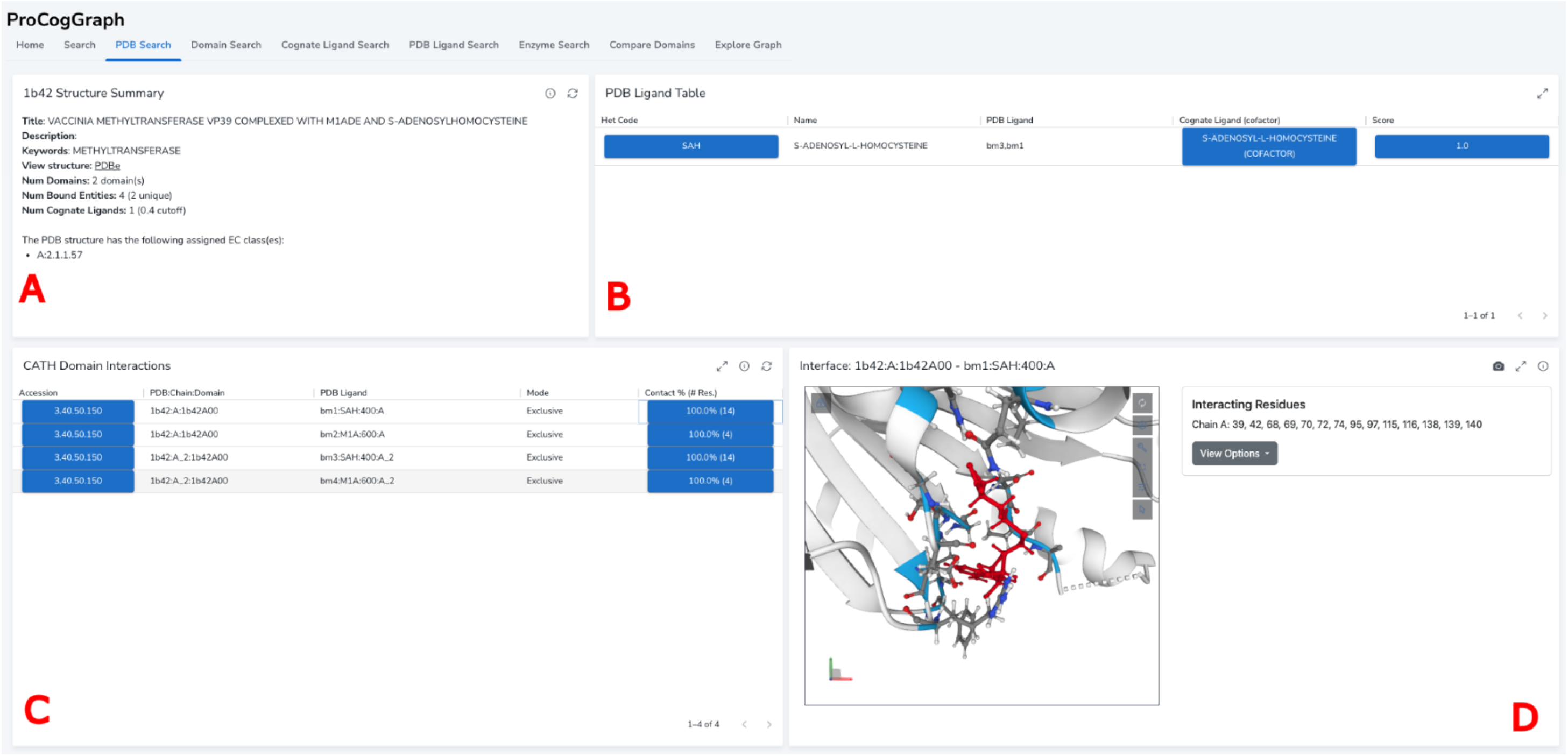
View of the ProCogGraph dashboard, PDB view. The PDB structure 1B42 has been selected from the search page, resulting in a summary reported being presented. A: Summary text is derived from the mmCIF file including title, description, and keywords if available. The number of domains present from the currently selected database (here CATH) is stated together with the number of unique bound entities in the structure. The current PARITY score cutoff is noted together with the number of cognate ligand matches found. B: The unique bound entities in the structure are listed, including Het Code, name and specific occurrences e.g. bound molecule 1 and 2. Clicking the het code for a ligand navigates and populates a summary report for the bound entity including structures it occurs in and a visualisation of it’s structure. The cognate ligand matches if available for a ligand are listed (click to view Cognate Ligand summary page) and the similarity score. Clicking the similarity score displays the two structures together with their MCS in the iFrame section of panel D. C: The domain interactions table lists each domain, the ligand it interacts with and the mode of interaction. Clicking on the domain sends users to a domain summary page, and clicking on the contact % column for a domain loads an embedded Molstar viewer display of the domain-ligand interaction. D: iframe display section – when clicking on a cognate ligand score, the viewer displays the structure of the compared ligands and the score between them together with the MCS highlighted. When clicking on a domain interaction, a Molstar viewer display of the interaction is shown, with interacting residues from the focussed domain highlighted in blue, all other interactions highlighted in purple.

## Supplementary Text

### 1. ProCogGraph Installation and Setup

#### Installation and Running the Database

##### Docker

ProCogGraph is both a pipeline for analysis of structures and a database of cognate ligand-domain mappings. To get started, the easiest method, described below, is to run ProCogGraph in a Docker container - for installation instructions for the database on bare metal, and for running the Nextflow pipeline see the [installation](docs/installation.md) guide.

1. Download and install Docker from the Docker website (https://www.docker.com/get-started)
2. Clone the ProCogGraph repository:

~~~
git clone m-crown/ProCogGraph
cd ProCogGraph
~~~
3. Run the setup script to download the latest flat files and create the necessary directories and Docker compose files if running on Linux/OSX:

~~~
./setup_docker_linux.sh
~~~ or for Windows (in Powershell with administrative access)

~~~
Set-ExecutionPolicy Unrestricted
./setup_docker_windows.ps1
~~~ This script creates the necessary directories for setting up the database, downloads the latest flat files from Zenodo and produces two yaml files on Linux/MACOS, and two powershell files on windows, one to build the database (run first time only) and one to run the database (run each time you want to start the database).
4. Run the build command: Linux/MACOS:

~~~
docker compose -f docker-compose-build.yml up
~~~ Windows:

~~~
./run_build.ps1
~~~
5. Run the database: Linux/MACOS:

~~~
docker compose -f docker-compose-run.yml up
~~~ Windows:

~~~
Set-ExecutionPolicy Unrestricted
./run_services.ps1
~~~ After running the Docker Compose script, three containers are started, one for the Neo4j database, one for the NeoDash dashboard and an Nginx server which serves the iframe visualisations available within the dashboard. The database can be accessed by navigating to ‘http://localhost:7474’ in a web browser to access the neo4j browser tool or connecting to ProCogDash via http://localhost:5005/. On linux, the compose-run.yml file can be modified (or the ‘run_services.ps1’ on Windows) to specify memory allocation for the Neo4j database, which can be adjusted as necessary for your system. Currently, these are not set by the install script, and so will operate with the memory configured in docker. To adjust these parameters add the following lines to the environment section of the compose_run.yaml file (or add as environment parameters in the ‘run_servcies.ps1’ file on Windows):
  – `NEO4J_server_memory_heap_initial__size=3600m`
  – `NEO4J_server_memory_heap_max__size=3600m`
  – `NEO4J_server_memory_pagecache_size=2g`
  – `NEO4J_server_jvm_additional=-XX:+ExitOnOutOfMemoryError`
6. Access the dashboard. The ProCogDash dashboard is built using NeoDash, a Neo4j plugin. The dashboard can be accessed by connecting to a running instance of the database in Docker at http://localhost:5005. The dashboard requires a username and password, which are set to ‘neo4j’ and ‘procoggraph’ by default.
7. To stop the database, run the following command: Linux/MACOS:

~~~
docker compose -f compose_run.yml down
~~~ Windows:

~~~
Set-ExecutionPolicy Unrestricted
./stop_services.ps1
Set-ExecutionPolicy Restricted
~~~

##### Neo4j

Installation instructions for running the database on bare metal, rather than Docker, are described below.

1. Download the latest database flat files from Zenodo https://zenodo.org/records/13165852/ and clone the ProCogGraph repository:

~~~
git clone m-crown/ProCogGraph
wget
~~~ https://zenodo.org/records/13165852/files/procoggraph_flat_files_v1-0.zip?download=1 `-O /path/to/database_flat_files`
2. Download and install Neo4j community edition from the https://neo4j.com/download/. The database was built using Neo4j version 5.
3. Copy the build script from the repository to the Neo4j database directory (e.g. neo4j-5.4.0) and the database flat files to the import directory:

~~~
     cp -r
/PATH/TO/PROCOGGRAPH_REPOSITORY/nextflow/bin/import_neo4j_data
.sh /PATH/TO/NEO4J_DATABASE/
     cp -r /PATH/TO/DATABASE_FLAT_FILES/*
/PATH/TO/NEO4J_DATABASE/import/
~~~
4. Run the build script:

~~~
cd /PATH/TO/NEO4J_DATABASE/
./import_neo4j_data.sh
~~~
5. Start the Neo4j database:

~~~
bin/neo4j start
~~~
6. Access the database by navigating to http://localhost:7474 in a web browser and update the default password (set to user ‘neo4j’ and password ‘neo4j’ by default).
7. Access ProCogDash via http://neodash.graphapp.io/. The dashboard can be loaded into Neodash by expanding the menu option in the bottom left of the screen, clicking the ‘+’ icon and importing the dashboard from a JSON file. Upload the file from the repository at ‘procogdash/dashboard.json’.

#### ProCogGraph Pipeline

The ProCogGraph pipeline is built using Nextflow for workflow management. To run the pipeline, follow these steps:

- The pipeline utilises data from a number of different sources to build the ProCogGraph database. To begin, prepare a data files directory using the table of data sources available in the ProCogGraph repository.
- Clone this repository and install dependencies:

~~~
   git clone m-crown/ProCogGraph
   cd ProCogGraph
   conda env create -f nextflow/envs/environment.yml
~~~
- Preprocess RHEA reaction files:

~~~
   cd /PATH/TO/DATA_DIR/
   python3 preprocess_rhea.py --rhea_ec_mapping rhea2ec.tsv -
-rhea_reaction_directions rhea-directions.tsv --rd_dir rd/ --
outdir. --chebi_names chebi_names.tsv.gz
~~~
- Produce final manifest file of structures to be processed:

~~~
    python3 download_mmcif.py --sifts_file
/PATH/TO/DATA_DIR/pdb_chain_enzyme.tsv.gz --assemblies_file
/PATH/TO/DATA_DIR/assemblies_data.csv.gz --chunk_size 100 --
output_dir /PATH/TO/STRUCTURES_DIR
~~~
- Run the nextflow pipeline. To configure the nextflow pipeline, the nextflow.config file within the repository should be modified. A SLURM cluster profile, specific for the development of the pipeline within the Bashton Group at Northumbria, is included called ‘crick’. The standard profile is designed for running the pipeline on a local machine, and is by default configured with a large amount of memory and CPU resources. This should be adjusted before running. Four additional parameters must be set specific to the user’s environment:
  - params.data_dir - the path to the data directory created including data files described above.
  - params.cache_in - the path to the cache directory for the pipeline, if pipeline has been run previously.
  - params.output_dir - the desired output directory.
  - params.manifest - the path to the manifest file created in step 3.

~~~
cd /PATH/TO/PROCOGGRAPH_REPOSITORY/nextflow nextflow
run main.nf -resume -profile standard
~~~

### 2. ProCogDash Implementation

The underlying ProCogGraph database can be explored using ProCogDash - a NeoDash dashboard which collects various sets of useful Cypher queries into a single source, running either locally or accessible from http://neodash.graphapp.io/. In contrast to past web versions of PROCOGNATE, similarity score cutoffs are user-customisable, from the minimum cutoff up to 1.0/perfect similarity. When exploring the range of ligands above the minimum cutoff, a user may want to search with a higher cutoff if multiple potential cognate ligands are identified for a PDB ligand, for example, to ensure docking studies utilise the most similar cognate ligand structure.

The dashboard contains five key visualisation modes as UI tabs: PDB, Cognate Ligand, PDB Ligand, Domain and EC. The homepage provides summary statistics for the number of structures and ligands represented in the current version of the graph and the number of cognate ligand matches for the currently specified cutoff. From the search page, global search settings can be specified, which consists of a similarity score cutoff for cognate ligand matching, and the domain database type to be searched against. A filter can also be applied to all cognate ligand matches to specify which cognate ligand matches should be considered:

- “All”, in which all bound entities for a PDB are displayed in the interaction table, regardless of whether they have a mapping to a cognate ligand.
- “Any”, in which all cognate ligand matches for a bound entity which are above the scoring threshold are presented.
- “Best” in which only cognate ligands with the highest score for a given bound entity are shown.

It should be noted that even when set to “Best”, a bound entity may have matches to multiple cognate ligands with the same maximum score. ProCogGraph is designed to serve as an information source, and so does not make an effort to select a particular best match as the “Best” best match, instead leaving this up to the user.

Next, users can initiate a search for one of each of the five visualisation modes. The PDB search box is used to search for a structure, and a PDB ID can be matched from any partial searches via a dropdown list. A clickable link is then presented next to the PDB search box, which takes you to the PDB visualisation mode.

On the PDB exploration page, a summary report is generated and presented, alongside a table of interactions between domains and bound entities in the structure, and the mapping of bound entities to cognate ligands.

Alongside the domain interaction table, an embedded iframe allows visualisation of the interacting residues between bound entities and domain residues in the structure, using PDBe-Molstar. In this visualisation, a list of interacting residues belonging to the selected domain are presented alongside the structure, with residues from the currently focussed domain highlighted in blue, and other domain residues highlighted in purple - this view is shown in Supplementary Figure 10.

The PARITY score between cognate ligands and bound entities can also be viewed, showing both the MCS match and the atom matches, making up the PARITY score - visualised with RDKit JS. An alternative to using RDKit JS for visualisation is the generation of image files for each structure. However, to view MCS structures and atom matches for all combinations of PDB and cognate ligands would necessitate a large amount of disk storage and significant computation. By dynamically generating these visualisations, the overall footprint of ProCogDash is minimised. The iframe visualisations are loaded from an Nginx server running in the distributed Docker image if running locally. Visualisation of the full contact annotation from PDBe-Arpeggio and SIFTS domain annotations is not possible with Molstar viewer, and so alongside this, ProCogGraph contains a script to generate Pymol sessions for each structure, displaying contacts and domain information on the assemblies using the information produced during pipeline execution.

For each domain listed in the PDB structure page, breakout links are accessible to a domain summary page (domains can also be searched for directly from the search page using the domain search box). This page includes a summary report of the number of ligands the domain is known to interact with, together with links to the external domain annotation. The report also summarises interactions for a domain at a “group” level, which varies depending on the domain database being examined; for example, in the CATH/Gene3D/SCOP/SUPERFAMILY, the group level is Superfamily, and for Pfam, it is the family level. Summaries are presented on the following:

- Group interactions table: lists all cognate ligands a group level are known to interact with in the database, together with the number of domains that interact with the ligand.
- Domain Contexts: This query lists the contexts in which a domain interacts with a ligand i.e, the other domains involved in the interaction and their interaction modes.
- Domain Cognate Ligand breakdowns: table lists the cognate ligands the specific domain searched for interacts with, and the percentage of the overall group the domain belongs to which also interact with the ligand. This is useful for identifying if the cognate ligand a domain binds to reflects typical superfamily activity or is an outlier.

Cognate and PDB ligands can be viewed in detail through breakout from the PDB structure page view. Both pages contain a similar set of results tables, including an RDKit.js visualisation of the ligand structure, a summary report detailing the cognate ligand database cross-references or the number of times a PDB ligand has been observed in the database, and the domain interactions that are observed for a ligand. Additionally, PDB ligands contain link tables to cognate ligand mappings. As with domains, PDB and cognate ligands can also be searched for directly from the search page, either by hetcode or name for PDBligands, or database ID (format DB:ID) or name for cognate ligand.

When searching for an EC number, results are aggregated and links presented to relevant structures, cognate and PDB ligands, together with a summary of domains known to interact wit ligands In this reaction. In addition the reactions associated with the EC number within the RHEA database are visualised using RDKit-js, allowing for dynamic generation based on the reaction smiles strings associated with the EC nodes in the graph.

The interlinked nature of the various results pages and their associated query results enable an intuitive and easy-to-use tool to drive analyses using ProCogGraph. An important aspect of any tool is the user-community, and one of the primary reasons for using Neo4J as a DBMS was the Neodash plugin which has been used to create ProCogDash. This plugin enables easy extensibility by users to create their own visualisations. Using a local version of the graph, which can be easily set up using the instructions provided in the repository, a user can connect to this graph in a few clicks and create a fresh dashboard that can be populated with their own research queries.

Every result presented in ProCogDash is generated using a Cypher query, which are contained within the neodash dashboard, and which is stored as a node within the database, allowing it to be versioned and distributed alongside the database itself. In addition to this, all queries are also made available within the repository as a single YAML file, where each query contains additional comments describing the underlying process.

Overall, the ProCogDash dashboard enables rapid and rich interrogation of the underlying database, in a highly adaptable and customisable manner due to the flexible Cypher queries underpinning the dashboard, which allow for high levels of filtering and customisation of the returned data depending on a user’s use case.

### 3. Common ligands with no cognate mapping

The most frequently observed bound entity descriptor with no cognate ligand match is Chlorophyll A (present in structures with EC reactions 1.97.1.12, 7.1.1.6 and 1.10.3.9). In these reactions, Chlorophyll A is referenced in the EC reaction comments text but is not present as a component of the reaction itself. Without inclusion in the reaction, ProCogGraph cannot assign this ligand as cognate, so remains unannotated. PC1, 3PE, PLX and PEK are all phospholipids, frequently found in liposomes and membranes, but not part of the enzyme reaction, hence they remain unmapped. Lipidic cubic phase crystallisation is the most commonly used method for crystallising membrane proteins, and varying mixes of all of these phospholipids have been studied for their effects on the structure of membranes formed in this crystallisation method (Caffrey 2015). Additionally, tetrofluoroaluminate and beryllium trifluoride ions are often used in combination with ADP/GDP to simulate binding of ATP/GTP binding sites in proteins (Bigay *et al*. 1987), which act as an analogue of the nucleotide triphosphate gamma phosphate (Pellegrini *et al*. 2024). In many cases, the metal fluoride is left unmatched whilst there is a cognate ligand match between ADP and cognate ATP *e*.*g*. in PDBs 1MMD, 4TYN and 3GLG.

## Notes

https://github.com/bashton-lab/ProCogGraph

https://doi.org/10.5281/zenodo.13165851

https://doi.org/10.5281/zenodo.13249841

### Competing Interest Statement

The authors have declared no competing interest.

